# Potential silencing of gene expression by PIWI-interacting RNAs (piRNAs) in somatic tissues in mollusk

**DOI:** 10.1101/2020.07.12.199877

**Authors:** Songqian Huang, Yuki Ichikawa, Kazutoshi Yoshitake, Yoji Igarashi, Mariom, Shigeharu Kinoshita, Md Asaduzzaman, Fumito Omori, Kaoru Maeyama, Kiyohito Nagai, Shugo Watabe, Shuichi Asakawa

## Abstract

PIWI/piRNA suppress transposon activity in animals, thereby safeguarding the genome from detrimental insertion mutagenesis. Recently, evidence revealed additional piRNA targets and functions in various animals. Although piRNAs are ubiquitously expressed in somatic tissues of the pearl oyster *Pinctada fucata*, their role is not well-characterized. Here, we report a PIWI/piRNA pathway, including piRNA biogenesis and piRNA-mediated gene regulation in *P. fucata*. A locked-nucleic-acid modified oligonucleotide (LNA-antagonist) was used to silence a single piRNA (piRNA0001) expression in *P. fucata*, which resulted in the differential expression of hundreds of endogenous genes. Target prediction analysis revealed that, following silencing, tens of endogenous genes were targeted by piRNA0001, including twelve up-regulated and nine down-regulated genes. Bioinformatic analyses suggested that different piRNA populations participate in the ping-pong amplification loop in a tissue-specific manner. These findings have improved our knowledge of the role of piRNA in mollusks, and provided evidence to understand the regulatory function of the PIWI/piRNA pathway on protein-coding genes outside of germline cells.

---

Animal species express three types of endogenous silencing-instigating small RNAs: microRNA (miRNA), endogenous siRNAs (endo-siRNAs), and PIWI-interacting RNAs (piRNAs), based on their biogenesis mechanism and type of Argonaute-binding partners^1^. miRNAs and endo-siRNAs are generated from double-stranded precursors by Dicer into fragments 20–23 nucleotides (nt) in length^1,2^. piRNAs are generated from single-stranded precursors in a manner independent of RNase III enzymes^3^, which are, by contrast, necessary for miRNA and endo-siRNA biogenesis. piRNAs associate with PIWI subfamily members of the Argonaute family of proteins, while miRNAs and endo-siRNAs associate with AGO subfamily members. The large Argonaute superfamily is characterized by the conserved PAZ domain, which is a single-strand nucleic acid binding motif^4^, and the PIWI domain, which implements RNase H slicing activity^5–7^.

The mechanisms underlying piRNA biogenesis and function remain largely unknown, mainly because the process has little in common with the miRNA and endo-siRNA pathways. However, remarkable progress has recently been made, especially in the area of piRNA biogenesis^8^. A comprehensive computational analysis of piRNA populations generated two models for piRNA biogenesis in various animals: the primary biogenesis pathway, and the amplification loop or ping-pong cycle^9,10^. In the primary biogenesis pathway, long piRNA precursors are transcribed from specific genomic loci called piRNA clusters, cleaved, and modified by intricate factors in the cytoplasm, before being transported into the nucleus by specific transcription factor complexes^10,11^. Primary piRNAs undergo an amplification process to induce high piRNA expression, known as the amplification loop or ping-pong cycle^9^. The Zucchini (Zuc) endonuclease potentially forms the piRNA 5’ end, while the 3’ end is 2’-O-methylted by HEN1/Pimet, associated with PIWI proteins^12,13^. Additionally, an uncharacterized 3’-5’ exonuclease such as Trimmer was observed to trim 3’ end of piRNAs in silkworms^14,15^. Hsp83/Shu may play a role in the PIWI loading step, and Hsp90/FKBP6 plays a role in secondary piRNA production^9^. Despite the recent characterization of several factors in the piRNA biogenesis pathway, it remains poorly understood.

PIWI proteins and their associated piRNAs suppress transposon activity in various animals, thereby safeguarding the genome from detrimental insertion mutagenesis^16–19^. However, many animals produce piRNAs that do not match transposon sequences determined in *Caenorhabditis elegans*^20^ and mouse genome^17^, suggesting additional piRNA targets and functions. Nucleic small RNA-mediated silencing has garnered great interest in recent years and numerous advances have been made in this field. Recent findings describing the role of transposon transcriptional regulation downstream of piRNAs have provided a paradigm for studying transcriptional regulation by small RNAs in animals, which, with the use of *C. elegans, Drosophila melanogaster*, and mice as simple model systems, can offer important new insights into this process^20,21^. For example, Saito et al postulated *in trans* regulation of the protein-coding transcript *Fas3* by piRNAs generated from the 3’ untranslated regions (3’UTR) of the *traffic jam* transcript^22^, while Robine et al determined that traffic jam protein levels were elevated in PIWI mutants, indicating a cis-regulatory mechanism for piRNA action in *D. melanogaster*^23^.

Mollusks are one of the largest groups of marine animals, of which *Pinctada fucata* is well-studied, owing to its economic potential for pearl production, as well as a model organism to investigate the fascinating biology of mollusks. Recent studies revealed that PIWI proteins are ubiquitously expressed in mollusks^24–26^. Moreover, abundant piRNA-sized small RNAs were observed in the oysters *Crassostrea gigas*^27,28^ and *Pinctada martensii*^29^, as well as in the scallop *Chlamys farreri*^30^. Our previous study characterized the ubiquitous expression of piRNAs in *P. fucata* somatic and gonad tissues^31^. However, piRNA biogenesis and their functions in *P. fucata* remain largely unknown. In the present study, our aims were to demonstrate the biogenesis and function of PIWI/piRNA, and investigate its endogenous gene regulation in *P. fucata*. Several key piRNA biogenesis factors (PIWI, Argonaute, Zucchini, and HEN1) were identified in *P. fucata*, and the primary and secondary biogenesis pathways were determined by comprehensive computational analysis. To understand piRNA-regulated endogenous gene expression, we examined the effect of silencing one of the most abundant piRNA (piRNA0001) on potential target genes such as miRNAs (predicted by a bioinformatic algorithm), using an LNA-antagonist.

## Results

### Transcriptomic analysis of piRNA biogenesis factors in *P. fucata*

Following clean-up and quality checks, sixteen transcriptomic libraries with 191.11 million reads, with a total length of 28.67 Gb, were used for *de novo* assembling (Table S1). A total of 324,779 transcripts were assembled, with a mean length of 564 bp (Table 1, RNA_Seq1). Complete sequences of PIWI (*Piwil1* and *Piwil2*), Argonaute (*AGO1* and *AGO2*), *Zucchini*, and *HEN1* were assembled based on the annotation results of transcripts, and their respective lengths are shown in Table 2. PIWI and AGO possessed the conserved PAZ and PIWI domains as homologs in other organisms, while the Zuc protein contained the PLDc domain (Fig. 1). The conserved identity of the PAZ and PIWI domains were 51.37% and 60.54%, respectively. Piwil1 and Piwil2 clustered with two categories of the PIWI subfamily, while AGO1 and AGO2 clustered with the vertebrate Argonaute subfamily and *Drosophila* AGO1 (Fig. S1).

**Table 1.**
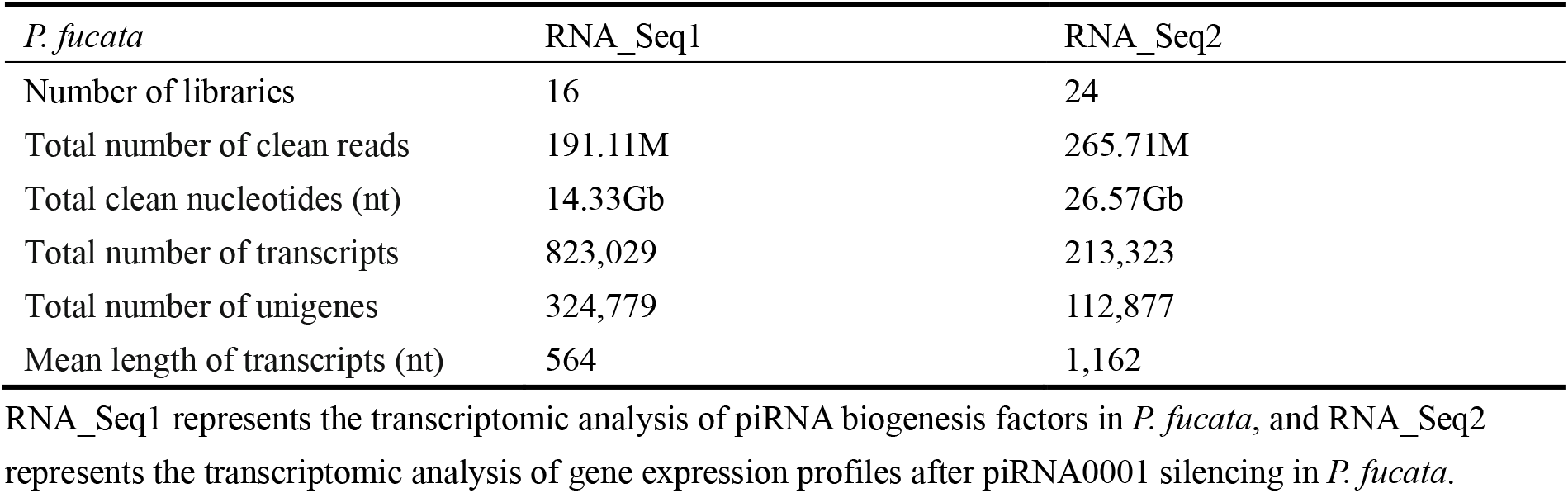
Summary of the mixed-assembling results of RNA sequencing in *P. fucata*.

| *P. fucata* | RNA_Seq1 | RNA_Seq2 |
| --- | --- | --- |
| Number of libraries | 16 | 24 |
| Total number of clean reads | 191.11M | 265.71M |
| Total clean nucleotides (nt) | 14.33Gb | 26.57Gb |
| Total number of transcripts | 823,029 | 213,323 |
| Total number of unigenes | 324,779 | 112,877 |
| Mean length of transcripts (nt) | 564 | 1,162 |

RNA\_Seq1 represents the transcriptomic analysis of piRNA biogenesis factors in *P. fucata*, and RNA\_Seq2 represents the transcriptomic analysis of gene expression profiles after piRNA0001 silencing in *P. fucata*.

**Table 2.**
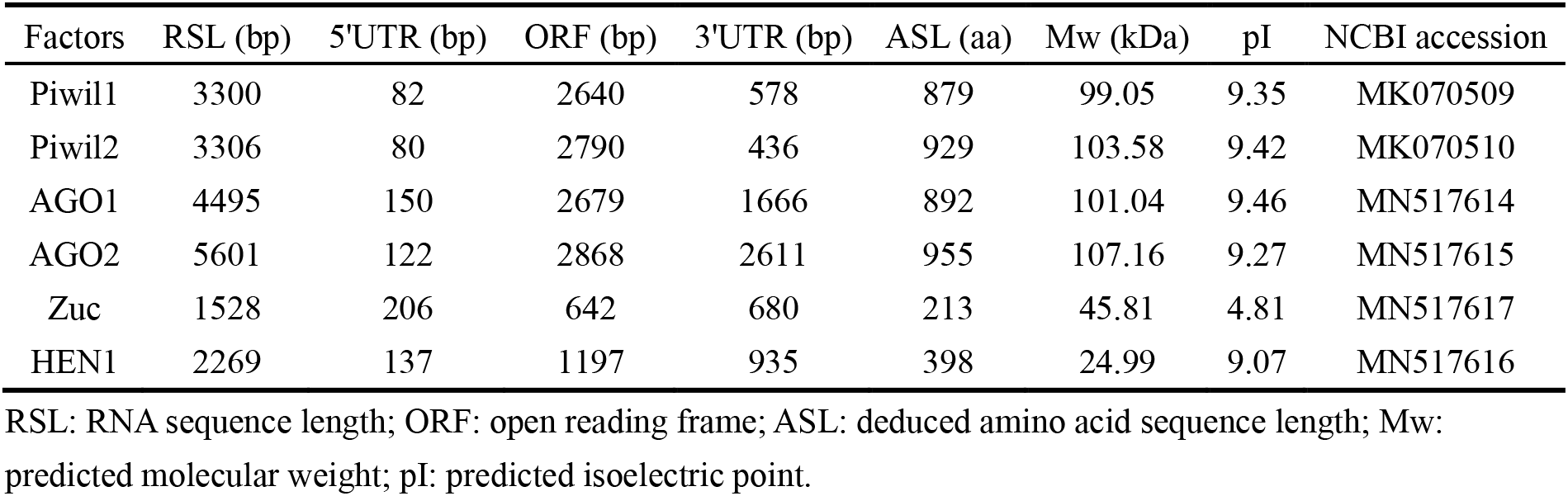
Assembling piRNA biogenesis factors in *P. fucata*.

**Figure 1.**
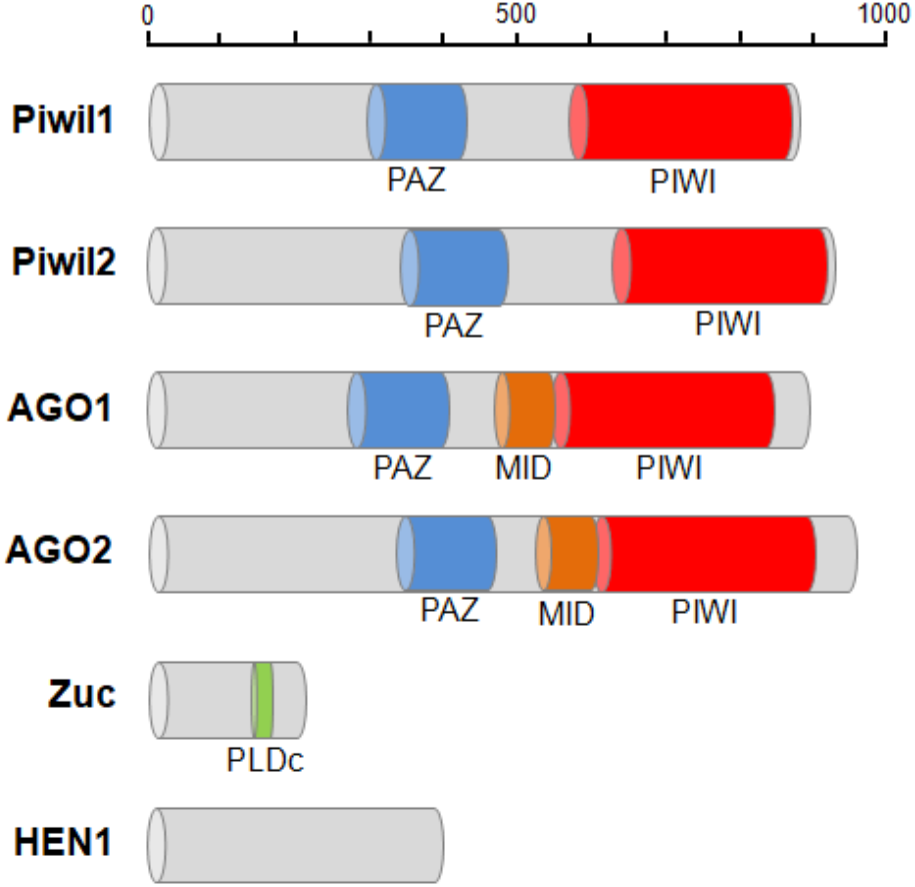
Protein domain structure of piRNA biogenesis factors in *P. fucata*. The PAZ domain contributes to specific and productive incorporation of miRNAs, endo-siRNAs, and piRNAs into the RNAi pathway; the PIWI domain can be inferred to cleave single-stranded RNA, e.g., mRNA, guided by double-stranded small RNAs; MID and PIWI domains place the guide uniquely in the proper position in Argonaute-RNA complexes. PLDc produces phosphatidic acid from phosphatidylcholine, which may be essential for the formation of certain types of transport vesicles or involved in constitutive vesicular transport via signal transduction pathways.

### Expression profiles of piRNA biogenesis factors in *P. fucata*

Gene expression was calculated as TPM by transcriptomic analysis. *Piwil1* and *Piwil2* were widely expressed in *P. fucata* somatic and gonad tissues, with a notably higher expression in gonad tissues at one- and three-year-old stages (Fig. 2a, b). *Piwil1* expression level was markedly higher than that of *Piwil2. AGO1* expression was varied and well balanced, while *AGO2* was highly expressed in female gonad tissues (Fig. 2c, d). *Zuc* was highly expressed in gonad tissues, especially in those of three-year-old females (Fig. 2e), and *HEN1* was expressed in all examined tissues (Fig. 2f). RT-PCR analysis validated *Piwil1* and *Piwil2* expression levels in larvae, juvenile, and adult *P. fucata*, which were high in fertilized eggs (0 hpf), sharply decreased in embryos (0.5 hpf), and weak from the blastula (4 hpf) to spat (30 dpf) stages (Fig. S2a, b). *Piwil1* and *Piwil2* were ubiquitously expressed in somatic and gonad tissues, and also displayed high expression levels in female *P. fucata*, both at the one- and three-year-old stages. While these piRNA biogenesis factors are generally highly expressed in the gonad tissues of other animals, they were weakly but ubiquitously expressed in somatic tissues, indicating their contribution to piRNA biogenesis in *P. fucata* somatic tissues.

**Figure 2.**
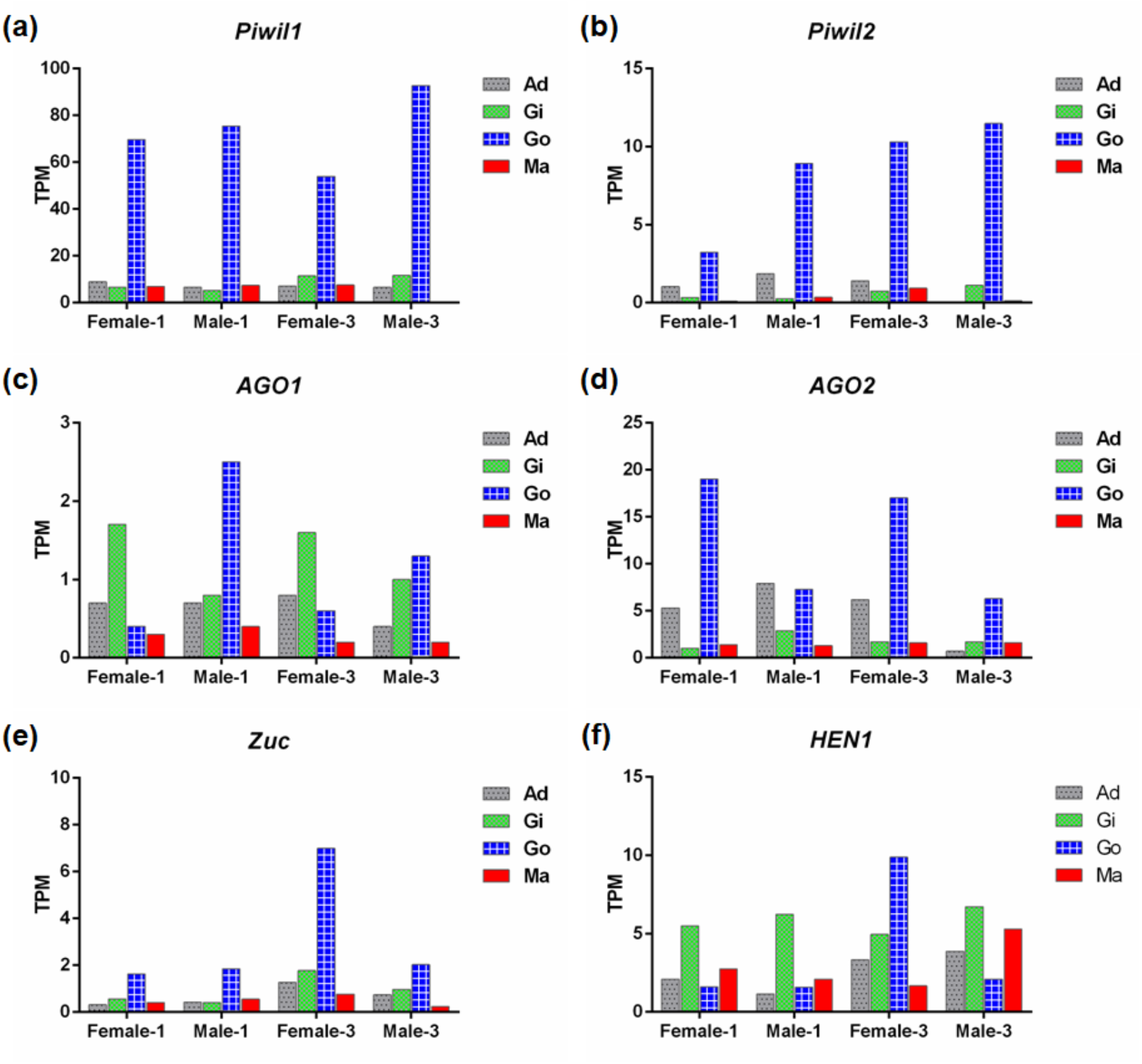
Expression abundance of piRNA biogenesis factors in *P. fucata* at different developmental stages, by RNA sequencing analysis. TPM: transcripts per million; Ad: adductor muscle; Gi: gill; Go: gonad; Ma: mantle.

### Ping-pong amplification loop of putative piRNAs in *P. fucata*

As *P. fucata* encodes two ubiquitously expressed PIWI proteins, we further studied the participation of distinct piRNA populations in the ping-pong amplification loop. We employed a bioinformatics approach, under the premise that Piwil1 and Piwil2 bind to piRNAs with different length profiles, similar to their corresponding mouse homologs Miwi and Mili, which preferentially bind to piRNAs 29/30 nt and 26/27 nt long, respectively^32^. We analyzed pairs of *P. fucata* sequenced reads with a 10-bp 5’ overlap (ping-pong pairs), which is the typical sequence length of each ping-pong partner (Fig. 3a and Fig. S3). In somatic tissues, ping-pong pairs combine piRNAs 29/30 nt in length, suggesting Piwil1-Piwil1-dependent homotypic ping-pong amplification. In gonad tissues, most ping-pong pairs combine piRNAs predominantly 29/30 nt and 25/26 nt in length, suggesting both Piwil1-Piwil2-dependent heterotypic and Piwil1-Piwil1-dependent homotypic ping-pong amplification, as depicted in Fig. 3b. Furthermore, 29/30 nt piRNAs that presumably bound to Piwil1 were heavily biased for a 5’ uridine (U), whereas 25/26 nt piRNAs that presumably bound to Piwil2 showed a stronger bias for an adenine (A) at position 10.

**Figure 3.**
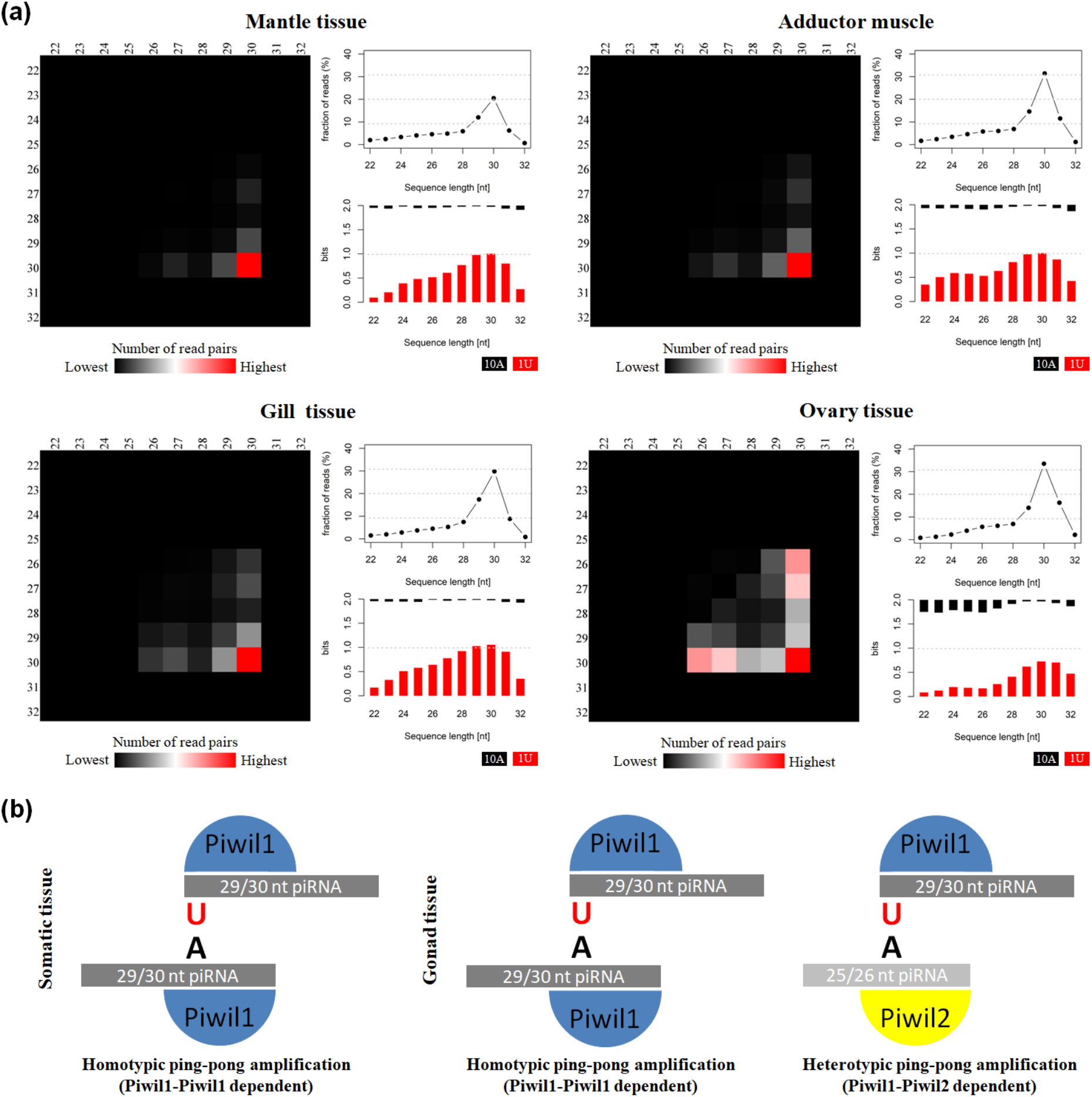
Analysis of putative piRNAs that participate in the ping-pong amplification loop in *P. fucata*. (a) Ping-pong matrices illustrate frequent length-combinations of ping-pong pairs (sequences with a 10-bp 5’ overlap). Sequence read length distribution and 1U/10A bias [bits] for ping-pong sequences are shown. (b) Proposed ping-pong amplification model in somatic (left) and gonad (right) tissues of *P. fucata*.

### piRNA0001 silencing in *P. fucata*

An LNA-antagonist was used to silence single piRNA (piRNA0001) expression in *P. fucata*. All samples were dissected for tissue collection two weeks after injection. Following RNA extraction, stem-loop RT-PCR was used for piRNA relative quantitative analysis. piRNA0001 expression levels in LNA-introduced pearl oysters were decreased in adductor muscle (0.04 ×, *p* < 0.01), gill (0.19 ×, *p* < 0.01), gonad (0.28 ×, *p* < 0.01), and mantle tissues (0.62 ×, *p* < 0.05), compared to that of the control group (Fig. 4a). Moreover, piRNA0001 precursor detected by transcriptomic analysis of gene expression profiles showed no significant differences in expression between LNA and Con groups (Fig. S4).

**Figure 4.**
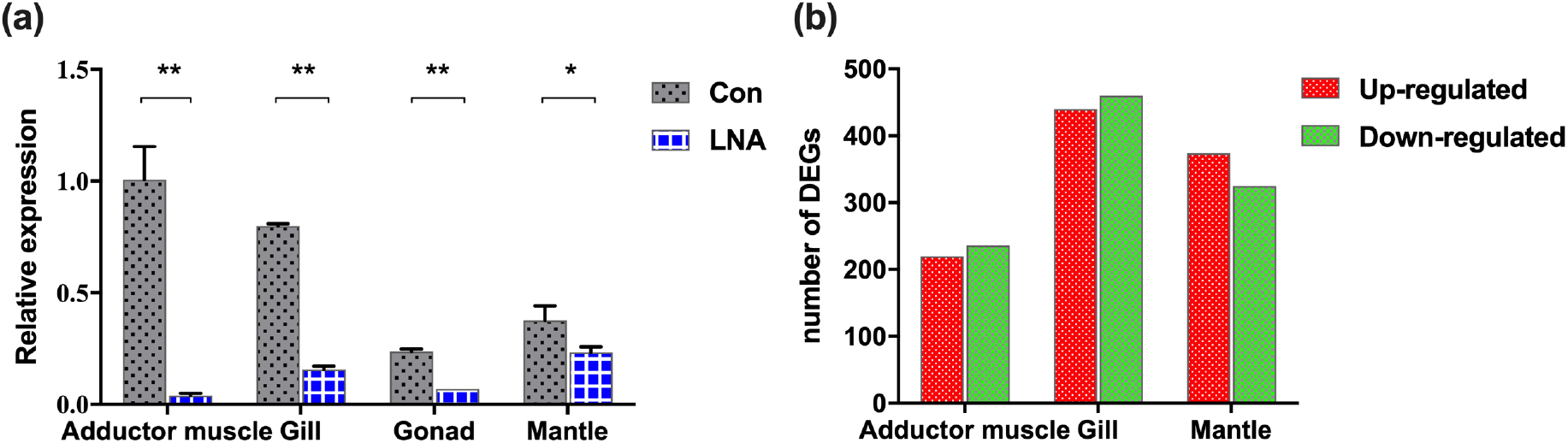
piRNA0001 silencing in *P. fucata*. (a) piRNA0001 expression profiles in *P. fucata* treated with LNA-antagonist. *U6* was used as the reference gene. The relative expression of piRNA0001 was normalized by piRNA0001 expression following LNA-antagonist treatment. Differences were statistically analyzed between Con and LNA groups using one-way analysis of variance (ANOVA). Significant differences are marked with * (*p* < 0.05) and ** (*p* < 0.01). (b) Number of differentially expressed genes (DEGs) between LNA and Con groups below the following threshold: *P*-value < 0.05 and folds > 2.

### Gene expression profiles following piRNA0001 silencing in *P. fucata*

Twenty-four libraries from Con and LNA groups, including adductor muscle, gill, gonad, and mantle tissues were constructed for RNA sequencing, with three replicates. Following data cleaning and quality checks, RNA sequencing produced 265.71 million clean reads (Table S2), i.e., a total length of 26.57 Gb, further assembled into 213,323 transcripts by Trinity (Table 1, RNA_Seq2). The Trinotate pipeline annotated 48.63% of these transcripts, based on five public databases (NT, Swiss-Prot, Pfam, GO and KEGG), and identified 31.06%, 63.37%, 80.60%, 60.88%, and 38.14% in the NT, Swiss-Prot, Pfam, GO, and KEGG database, respectively. Additionally, most of these transcripts were blasted with *P. fucata* and other mollusks species (Fig. S5).

Analysis of differentially expressed genes (DEGs) in somatic tissues between Con and LNA groups was carried out to further understand the molecular events involved in the functions of piRNA0001 in *P. fucata*. We selected the genes from at least one group with a mean TPM value > 5 for analysis, using edgeR (*P*-value < 0.05 and folds > 2). A total of 2,904 transcripts showed significant differential expression between Con and LNA groups (Table S3). Pairwise comparison in adductor muscle revealed 456 differentially expressed transcripts, including 220 up- and 236 down-regulated after LNA-antagonist treatment (in comparison with the Con group).

Additionally, analysis of 900 transcripts revealed 440 up- and 460 down-regulated DEGs in gill tissue, as well as 374 up- and 325 down-regulated DEGs in mantle tissues, following LNA-antagonist treatment (Fig. 4b). Among these, 16 and 15 DEGs were typically up- and down-regulated in the three examined LNA-antagonist-treated tissue types, respectively. All DEGs retrieved from the comparison between Con and LNA groups were used for function enrichment and classification analysis. The most significantly enriched categories were cellular process (GO:0009987) and single-organism process (GO:0044699) in biological process, and cell (GO:0005623), cell part (GO:0044464) and organelle (GO:0043226) in cellular component, and binding (GO:0005488) and catalytic activity (GO:0003824) in molecular function (Fig. S6). Furthermore, 78 KEGG pathways in the pearl oyster were enriched (*p* < 0.01) following piRNA0001 silencing (Fig. S7).

### Prediction of piRNA0001 targeting sites in *P. fucata*

All DEGs were used for piRNA0001 target prediction, with genes containing target sites in 3’UTR considered as potential target genes. Four, nine, and eleven DEGs were predicted to be targeted by piRNA0001 in adductor muscle, gill, and mantle tissues, respectively, with an average alignment score of 162.17 and average energy value of −14.00 kcal mol^−1^ (Fig. 5). Target prediction was anticipated for *FHOD3, SRS10, VINC*, and *ZNF622* in adductor muscle, for *ART2, EF1A, EMC7, ESI1L, NAC2, PA2HB, SYWC, TM87A*, and *ZNF622* in gill tissue, and for *CAH14, CBF, CDC42, FRRS1, GOLI4, KAT3, NBR1, PA2HB, TYR1, TYR2*, and *ZNF622* in mantle tissue. The predicted piRNA:mRNA interaction is shown in Table 3, including two sites on both the *ART2* and *CBF* 3’UTR. No polyA signal was observed in the assembly transcripts of *ART2, EF1A, ESI1L, FHOD3, PA2HB, SRS10*, and *TM87A*, and predicted interaction sites on *CAH14, NBR1, SYWC* and *VINC* were located after the polyA signal (Supplementary sequences). The second piRNA:mRNA interaction site on *ART2*, as well as interaction sites on *NAC2* and *VINC*, were located far from the stop codon (Table 3). Alignment pairing from the second to eighth nucleotides, considered to be the piRNA seed region, showed prefect complementarity. Furthermore, base-pairing outside of the seed region was also important for piRNA target prediction. Following piRNA0001 silencing in *P. fucata*, *FHOD3, SRS10*, and *ZNF622* were up-regulated, while *VINC* were down-regulated in adductor muscle tissue. In gill tissue, *ART2, EF1A, EMC7, ESI1L, SYWC, TM87A*, and *ZNF622* were up-regulated, while *NAC2* and *PA2HB* were down-regulated. In mantle tissue, *CBF*, *CDC42, NBR1*, and *ZNF622* were up-regulated, while *CAH14, FRRS1, GOLI4, KAT3, PA2HB, YTR1*, and *TYR2* were down-regulated. *ZNF622* was up-regulated in all examined tissues. Up- and down-regulated genes were both predicted to be targeted by piRNA0001, demonstrating the diverse ways of piRNA-mediated gene regulation in *P. fucata*.

**Table 3.**
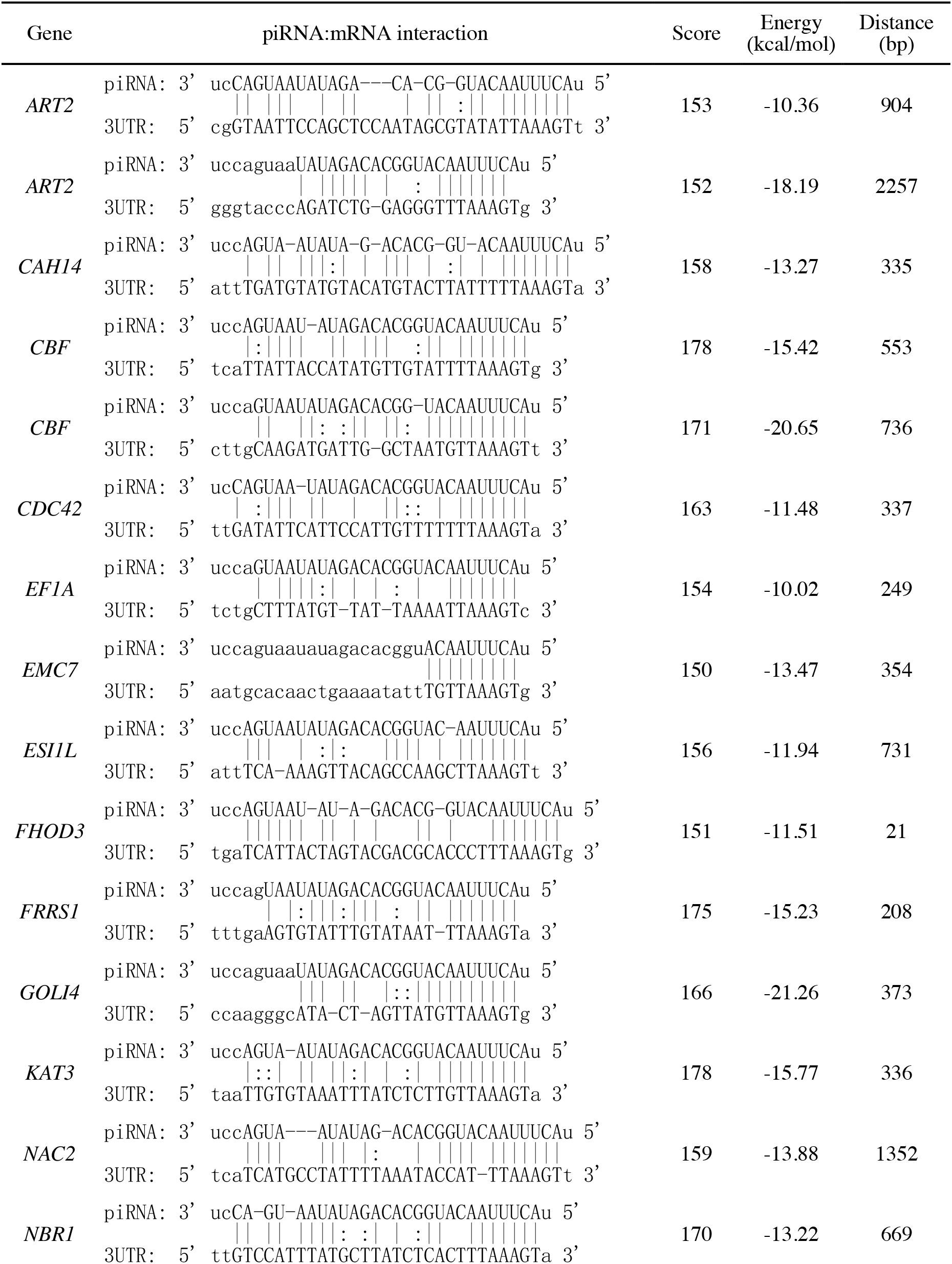
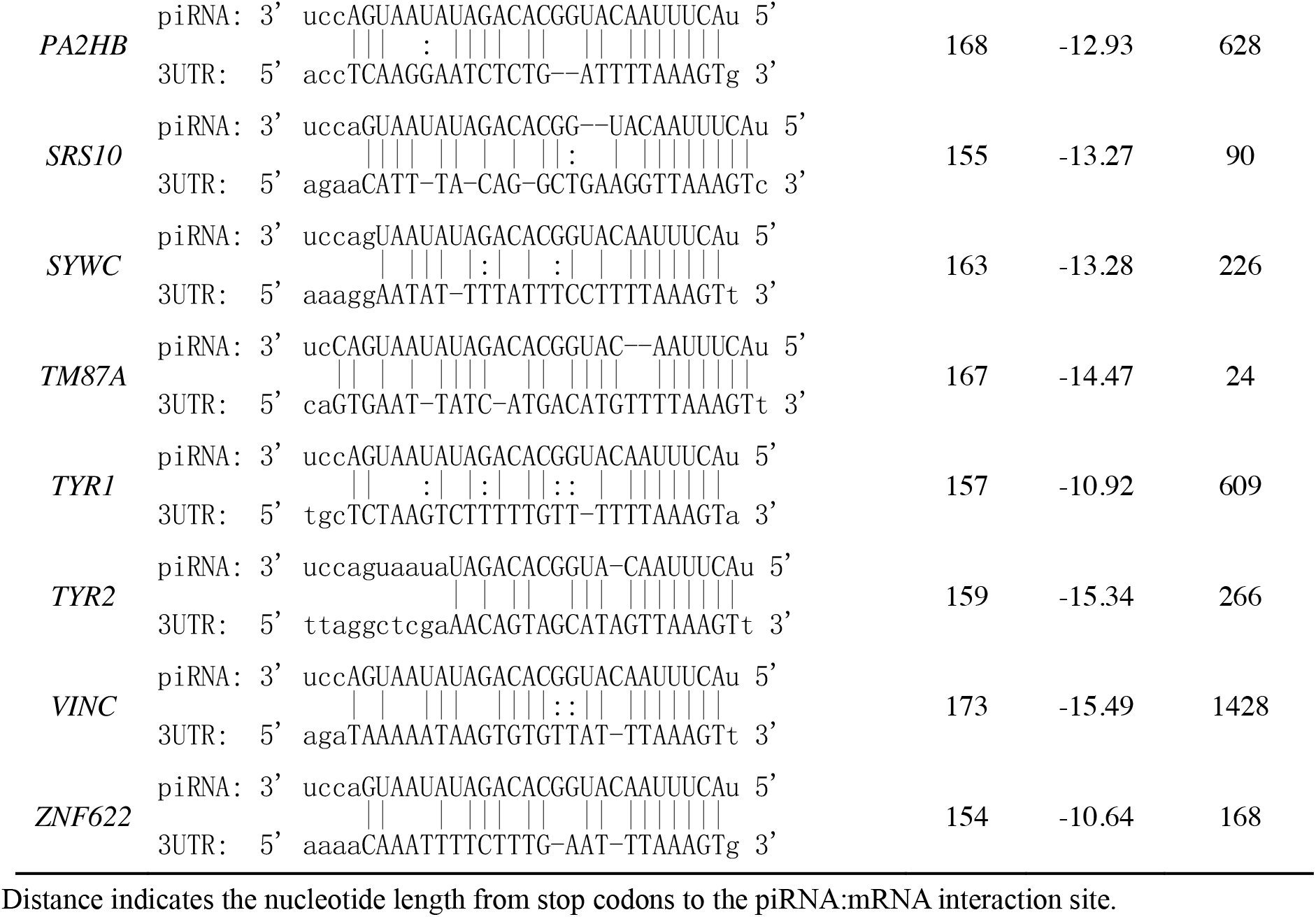
Predicted piRNA:mRNA interaction in *P. fucata*.

**Figure 5.**
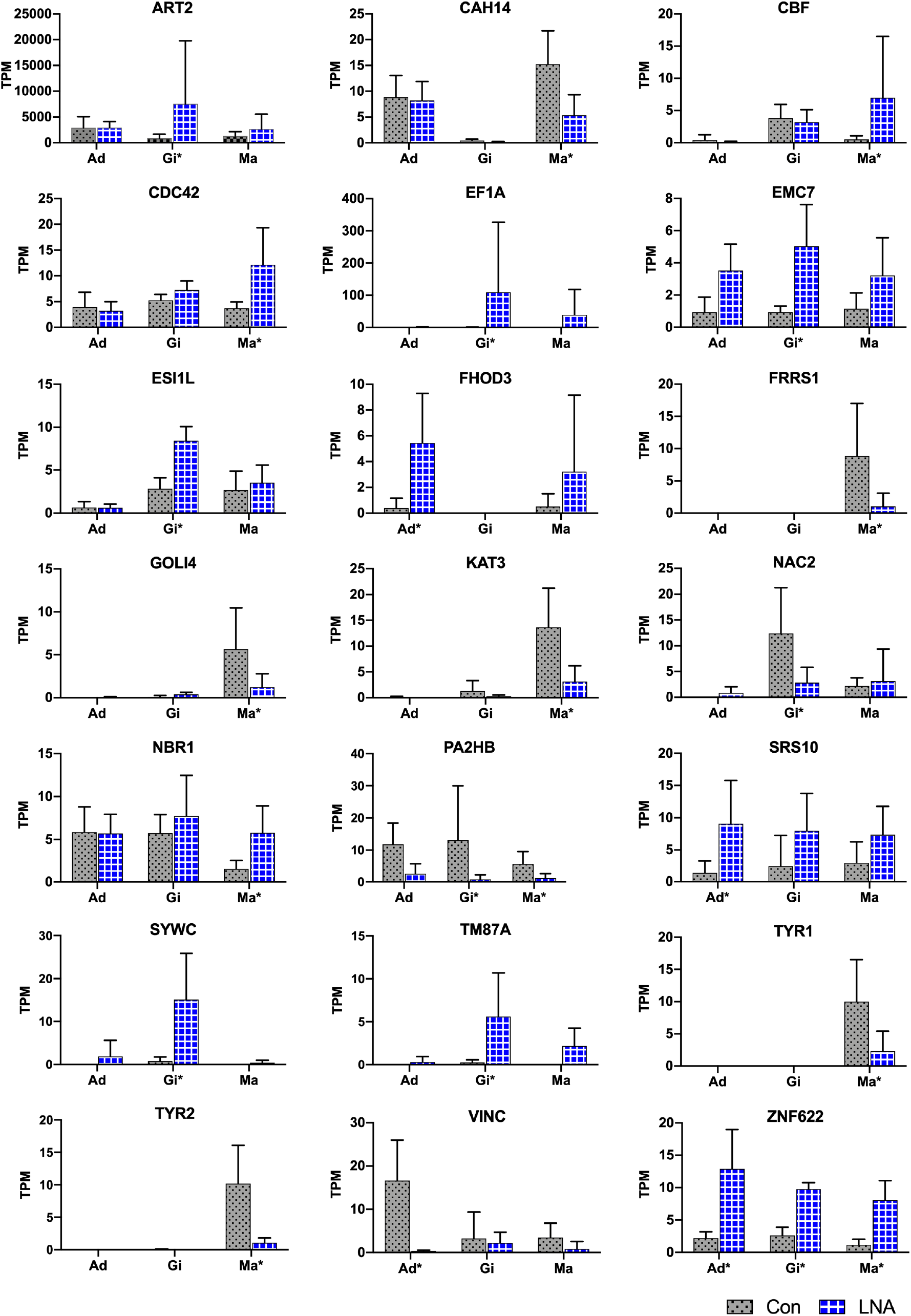
Target predictions of DEGs in *P. fucata*. TPM shows the relative expression level of target genes by RNA-seq. Tissue: adductor muscle (Ad), gill (Gi), and mantle (Ma). Tissue marked with an asterisk (*) represents differentially expressed gene under the following threshold: *P*-value < 0.05 and folds > 2 in the present tissue.

### Validation of gene expression by RT-PCR

Total RNA extracted from LNA and Con *P. fucata* somatic tissues was analyzed using RT-PCR, to determine the authenticity of mRNA expression calculated by RNA sequencing. An inconsistency was detected in adductor muscle *VINC* expression obtained from RNA sequencing and RT-PCR analysis (Fig. 6), possibly caused by ineffective RT-PCR primers or by sequencing errors that occurred during the complex process, with the predicted products located after the polyA signal on the 3’UTR, far from the stop codon (Supplementary sequences: *VINC*). Nonetheless, the remaining eight predicted target genes demonstrated consistent levels of relative expression via RT-PCR, compared with results obtained from RNA sequencing. Among these genes, *ZNF622* was up-regulated in all examined somatic tissues, following piRNA0001 silencing in *P. fucata*, while *PA2HB* and *TYR1* were down-regulated in gill and mantle tissues, both based on data obtained from RNA sequencing and RT-PCR.

**Figure 6.**
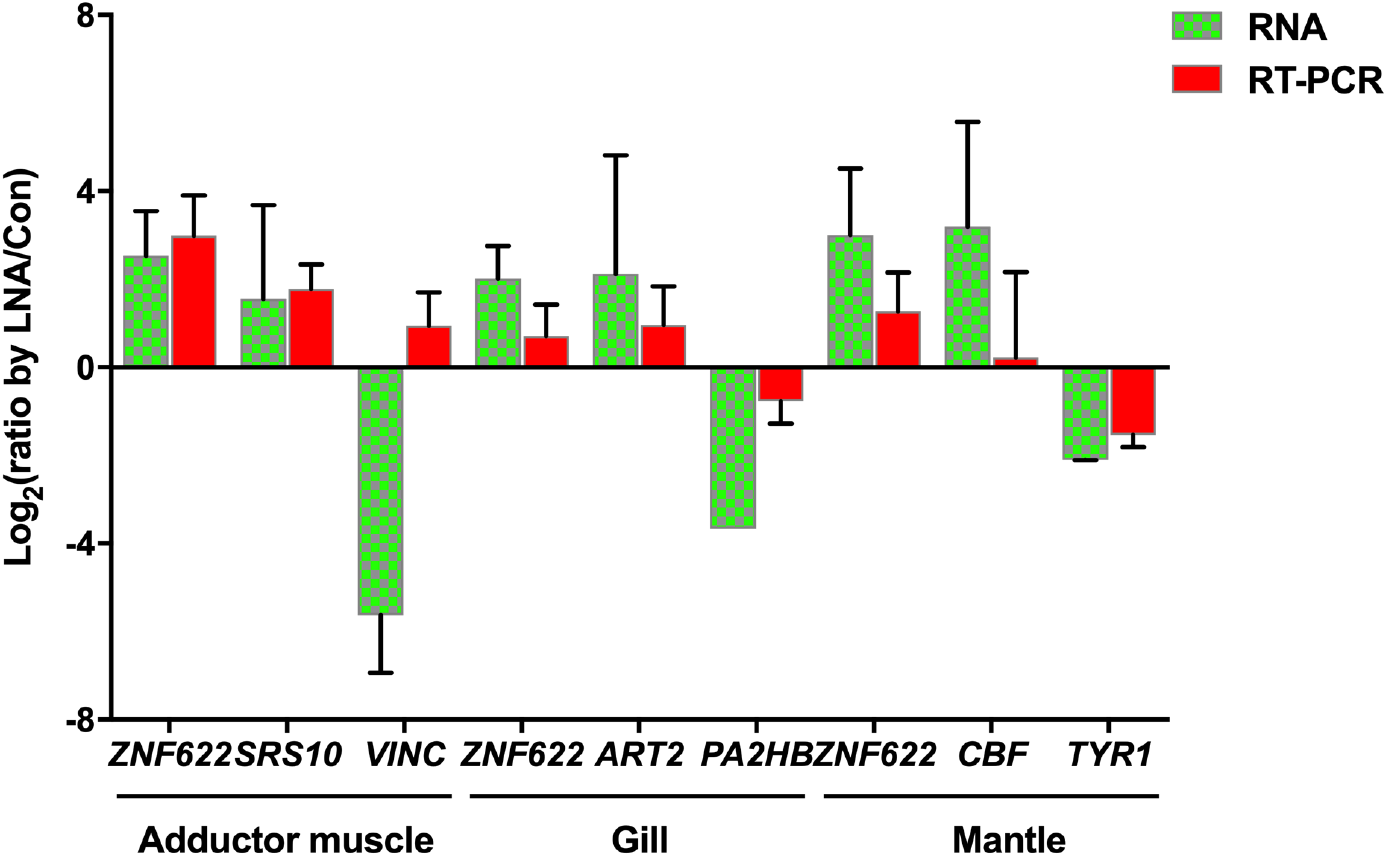
Validation of potential piRNA0001 predicted target genes in *P. fucata*.

## Discussion

Analysis of the PIWI/piRNA pathway representative of several animals revealed an extensive diversity of lineage-specific adaptations, challenging the universal validity of insights obtained from model organisms. The PIWI family belongs to a large subfamily of Argonaute proteins, which represent a large protein family in archaebacteria and eukaryotes^33^. These proteins are at the core of an RNA silencing machinery that uses small RNA molecules as guides to identify homologous sequences in RNA or DNA^3^. Argonaute proteins are characterized by two protein domains, namely the PAZ domain, an RNA-binding motif that binds the 3’ end of short RNAs, and the PIWI domain, which is structurally similar to the RNaseH catalytic domain^34^. In animals, Argonaute proteins can be further subdivided into two distinct classes: AGO proteins, which function together with siRNAs and miRNAs, and PIWI proteins, which associate with piRNAs in mammals. Four homologous *Hiwi, Hili, Hiwi2*, and *Hiwi3* were identified in *Homo sapiens*, and play a crucial role in human cancer and male germline cell development^35–37^. The homologous *Miwi, Mili*, and *Miwi2* were identified in *Mus musculus*, and their knockdown lead to male sterility^38–40^; Two homologues, Ziwi and Zili, were identified in *Danio rerio*, and were both crucial for germ cell differentiation and meiosis^3,41^. These findings indicated that PIWI may play an important role in germline cell development in organisms.

As PIWI was originally discovered in *Drosophila*, where it functions in germline cell maintenance and self-renewal^42^, PIWI/piRNA research in germline cells has rapidly advanced. PIWI mutations lead to a profound infertility phenotype in mice^18,38^. In addition to PIWI function in germline cells, new discoveries demonstrated the expression and function of PIWI in the soma, from low eukaryotes to mammals. PIWI expression was detected in human cancer cells^10,43^ and in mammalian somatic tissues^44^. Although somatic PIWI was previously detected in *Drosophila* fat body and ovarian somatic tissues^45,46^, further evidence of somatic PIWI/piRNA expression was observed in 16 out of 20 surveyed arthropod species^47^. In mollusks, PIWI proteins were widely expressed in *Lymnaea stagnalis* and *C. gigas* somatic tissues, and homologous PIWIs were also detected in 11 mollusks species, suggesting the occurrence of somatic PIWI/piRNA expression in early bilaterian ancestors^26^. In the present study, *Piwil1* and *Piwil2*, which were assembled by RNA sequencing in *P. fucata*, were ubiquitously expressed in all examined somatic tissues as well as gonad tissues. These results provide further evidence of somatic PIWI/piRNA expression as an ancestral bilaterian trait^47^.

piRNA biogenesis in metazoa involved the synthesis of primary piRNA from a piRNA cluster, assisted by several factors in the cytoplasm and nucleus. The long, single-stranded piRNA precursor was then exported from the nucleus (Fig. 7) to the cytoplasm, where it was shortened into piRNA-like small RNAs by an undetermined endonuclease^9,10^ (Ishizu et al., 2012; Ross et al., 2014). Recent studies have indicated that Zuc may be the endonuclease forming the 5’ end of piRNAs in *Drosophila* and mice^8,13,48^. The 3’ end was 2-O-methylated by HEN1^12^, while an uncharacterized 3’-5’ exonuclease was shown to trim the 3’ end of piRNAs^14,15^. In the present study, several crucial piRNA biogenesis factors, including PIWI, AGO, Zuc, and HEN1, were assembled by RNA sequencing. These were ubiquitously expressed in all examined tissues, especially in gonads, which were previously shown to exhibit a higher percentage of piRNA expression than somatic tissues^31^. The homogenous AGO1 expression may be owing to ubiquitous miRNA biogenesis in tissues, while the highly expressed AGO2 in gonad tissues may also play a crucial role in piRNA biogenesis in *P. fucata*.

**Figure 7.**
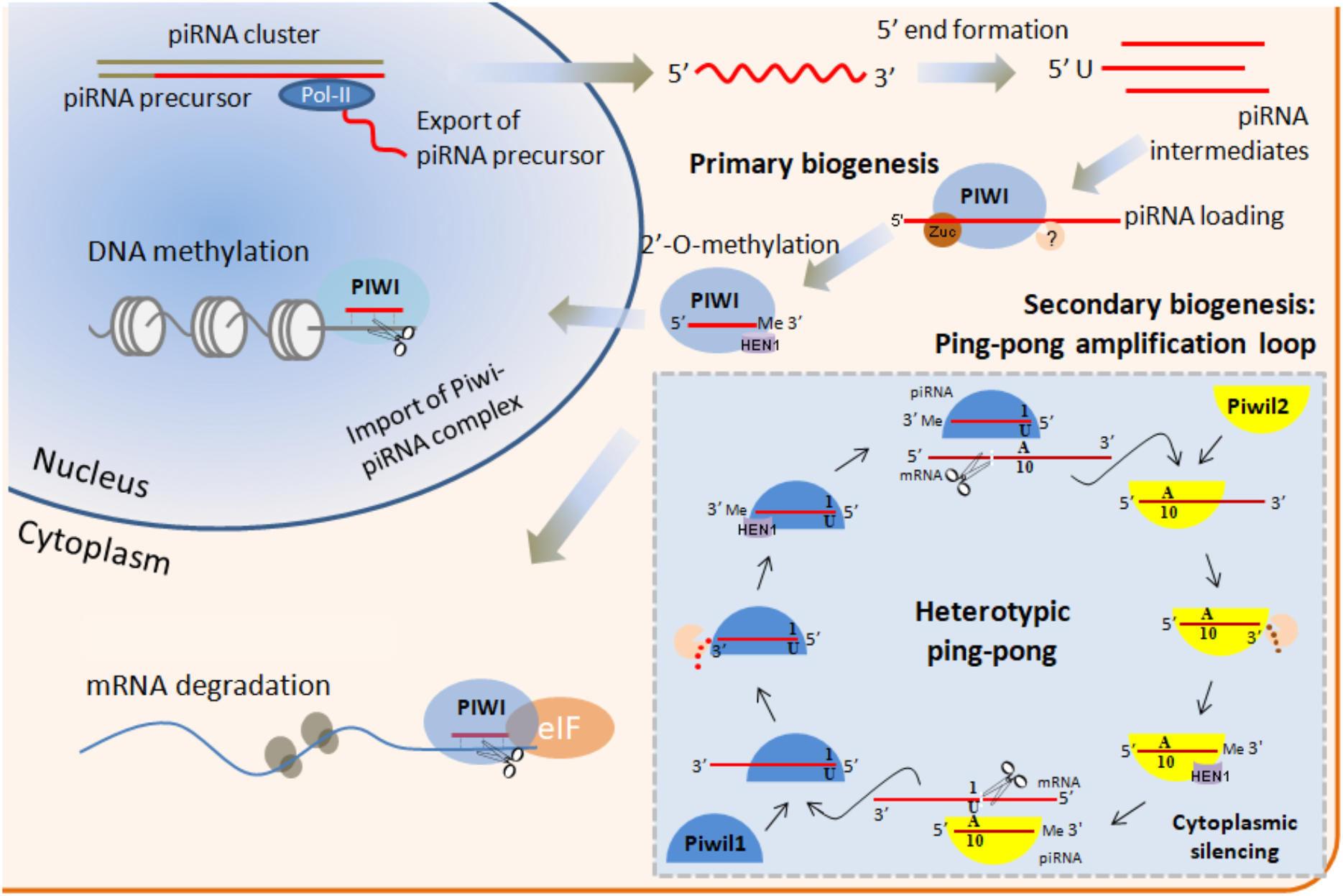
piRNA biogenesis in *P. fucata* consists of the primary piRNA processing pathway and the amplification loop. The primary transcripts of piRNA clusters are shortened into piRNA intermediates, which are loaded onto PIWI proteins. They are then trimmed from the 3’ end to the size of mature piRNAs, and 2’-O-methylated. PIWI associated with piRNAs is translocated to the nucleus. Piwil1 associated with primary piRNAs (blue) contributes to the ping-pong amplification loop to produce piRNAs that associated with Piwil2 (yellow) in gonad tissues. Piwil1/Piwil1 may operate a homotypic ping-pong amplification in somatic tissues, while the contribution of Piwil2 to this amplification loop may be negligible.

The ping-pong amplification loop is responsible for the post-transcriptional silencing of transposable elements^26^. In *Drosophila* and mice, this process typically involves two PIWI proteins (heterotypic ping-pong), one loaded with antisense piRNAs targeting piRNA cluster transcripts, which contain transposon sequences in antisense orientation^49^. The homotypic Aub:Aub ping-pong process occurred in *Drosophila*^50^ and wild-type prenatal mouse testes (Miwi2:Miwi2, Mili:Mili)^49^. In *P. fucata*, we also determined the amplification system for a secondary piRNA biogenesis pathway using comprehensive computational analysis. A Piwil1-Piwil1-dependent homotypic ping-pong amplification loop was observed in somatic and gonad tissues, with a Piwil1-Piwil2 dependent heterotypic ping-pong amplification loop simultaneously occurring in gonad tissues Nonetheless, we clearly cannot rule out the possibility that binding preferences of PIWI proteins have changed in *P. fucata*. One could even speculate that both PIWI proteins may bind the whole range of piRNAs. However, based on the presence of piRNA populations with length profile lengths and their representation in ping-pong pairs, together with the differences in their number of 1U and 10A reads, we believe the above explanation to be a reasonable and parsimonious interpretation of the data, while acknowledging the possibility of others.

The expression of piRNA biogenesis factors in somatic tissues implies the potential existence of somatic piRNAs. Recent studies have revealed the somatic piRNA from sponges to humans^10,44^. A functional non-gonadal somatic piRNA pathway in *Drosophila* fat body affected normal metabolism and overall organismal health^46^. Somatic piRNAs were considered an ancestral trait of arthropods, which predominantly targeted transposable elements, suggesting that the piRNA pathway was active in the soma of the last common ancestor of arthropods to keep mobile genetic elements in check^47^. Abundant putative piNRAs were observed in somatic and gonad tissues of *P. fucata* in our previous study^31^.

To explore the function of piRNA in *P. fucata*, LNA-antagonist was used to silence single piRNA (piRNA0001) expression. LNA-antagonist has been used for mouse and non-human primate small RNA silencing without affecting health (Elmén et al., 2008). In the present study, we used an LNA-antagonist to silence piRNA0001expression, which was the most highly expressed piRNA in *P. fucata* somatic tissues, followed by stem-loop RT-PCR for piRNA0001 quantification analysis, as in previous studies^20,51,52^. Stem-loop RT-PCR is a powerful and reliable tool for quantitatively analyzing and monitoring dynamic changes in piRNAs, specifically at the cellular and tissue levels in invertebrates. Here, piRNA0001 was effectively down-regulated in somatic tissues by the specific LNA-antagonist.

Recent studies indicated that piRNAs may regulate endogenous mRNA expression in *Drosophila* and mice^53,54^. Although targeting rules were determined in *Drosophila*^20,54^, the available tools cannot be used for piRNA targeting site prediction in other species. An RNA-RNA interacting prediction tool, such as miRanda^53^, was alternatively used to predict potential targeting sites between piRNA and endogenous genes. Here, we used miRanda to predict potential piRNA:mRNA interaction sites on differently expressed genes in *P. fucata*. piRNA target recognition was strikingly similar to that of miRNA pairing in the seed sequence (positions 2–8 relative to the 5’ end of the piRNA), which is the primary determinant of target recognition. Seed pairing is therefore not sufficient by itself, and additional base-pairing outside of the seed sequence is also important for piRNA target sites.

Among the predicted piRNA0001 target genes in *P. fucata*, the majority were related to metabolism and biological processes. *ZNF622* was up-regulated in all examined tissues following piRNA0001 silencing, and was predicted to be targeted by piRNA0001 with an alignment score of 154.00 and energy value of −10.64 kcal·mol^−1^. Zinc finger proteins (ZNFs) are one of the most abundant groups of proteins, with a wide range of molecular function, and are implicated in transcriptional regulation, ubiquitin-mediated protein degradation, signal transduction, actin targeting, DNA repair, cell migration, and numerous other process^55^. A single piRNA (*fem* piRNA) from a sex chromosome downregulated *Masc* mRNA, which encodes a C3H-type zinc finger protein that induces masculinization and plays an important role in sex determination in silkworms^56^. *CAH14, NBR1, SYWC*, and *VINC* were predicted targets of piRNA0001, while interaction sites were located after the polyA signal on the 3’UTR, suggesting that these interaction sites may be false positive results. Alternatively, this could have been caused by the insufficient assembly result of RNA sequencing, with the polyA signal not appearing in the assembling gene sequences, as observed in *ART2, EF1A, ESI1L, FHOD3, PA2HB*, *SRS10*, and *TM87A*. Single and double target sites were both observed in these target genes, indicating that piRNA may regulate endogenous genes by multiple sites on the 3’UTR. Except for up-regulated genes predicted to be targeted by piRNA0001, nine down-regulated genes may also have been targeted by piRNA0001, extending the uncertainty surrounding piRNAs in *P. fucata*. The lowest piRNA silencing efficiency was observed in mantle tissues, possibly explaining their majority of down-regulated genes. Although, we cannot finalize the target rules or genes of piRNA, we have provided new insights into the gene regulatory function of piRNAs in *P. fucata*. The PIWI/piRNA might have a great diversity of functions in *P. fucata*, including but not limited to gene regulation. Further research is needed to fully understand somatic piRNAs in mollusks.

In conclusion, piRNA biogenesis factors, including PIWI, AGO, Zuc, and HEN1, were assembled by RNA sequencing, and were ubiquitously expressed in all examined tissues. A homotypic and heterotypic ping-pong amplification system in *P. fucata* may also contribute to secondary piRNA production in a tissue-specific manner. Transcriptomic analysis of somatic tissue gene expression indicated that piRNA0001 may play a crucial role in endogenous gene regulation in *P. fucata*. These findings have successfully contributed to our understanding of the role of piRNA in mollusks and will also help gain further insight into the PIWI/piRNA pathway function outside of germline cells.

## Methods

### Transcriptomic analysis of piRNA biogenesis factors in *P. fucata*

Six (female: male = 1: 1) one-year-old and six (female: male = 1: 1) three-year-old pearl oysters were collected from the Mikimoto Pearl Research Institution Base, Mie Prefecture, Japan in May 2018. Adductor muscle (Ad), gill (Gi), mantle (Ma) and gonad (Go) tissues were collected and individually transferred to 2-mL tubes containing RNA later (QIAGEN, Maryland, USA) separately. The samples were stored overnight at 4 °C, and preserved at −80 °C until further analysis. Total RNA was isolated from each sample using the RNeasy Mini Kit (QIAGEN), according to the manufacturer’s instructions. After assessing RNA quality and quantity using the Agilent 2200 TapeStation (Agilent Technologies, Waldbronn, Germany), three isolated total RNA samples were mixed at equivalent concentrations to construct an RNA sequencing library for each tissue sample at different ages. A total of 2 μg mixed total RNA was used for library construction according to the manufacturer’s protocol, and subjected to paired-end sequencing on the Illumina HiSeq 4000 platform. High quality reads were required for *de novo* assembly analysis. Before assembly, raw reads were trimmed by removing adapter sequences and low-quality reads. Clean reads were jointly assembled into unigenes using the Trinity software^57^, and finally, transcriptome annotation was performed using the Trinotate pipeline (https://trinotate.github.io/)^58^.

The consensus sequences of piRNA biogenesis factors were assembled using CLC Genomics Workbench version 8.0.1 (QIAGEN) and confirmed by published homolog sequences using BLASTp. Open reading frames (ORFs) were identified using the ORF finder available at NCBI (https://www.ncbi.nlm.nih.gov/orffinder/). Structural features of deduced protein sequences were analyzed via SMART (http://smart.embl-heidelberg.de/)^59^. Predicted molecular weights and isoelectric points were determined using ExPASy^60^. All nucleotide and deduced amino acid sequences were edited electronically using BioEdit^61^. Sequence similarity of deduced proteins was performed using BLAST (https://blast.ncbi.nlm.nih.gov/Blast.cgi). Homologous sequences of piRNA biogenesis factors were downloaded from NCBI Protein databases and the alignment of amino acid sequences was performed by MEGA5.0^62^. The Neighbor-Joining method from MEGA 5.0 was used for building phylogenic trees with 1000 bootstrap replications.

### Ping-pong amplification loop of piRNAs biogenesis in *P. fucata*

Eight small RNA libraries from *P. fucata* adductor muscle, gill, ovary, and mantle tissues were sequenced by the Ion Proton system in our laboratory (the sequencing raw data were stored in the DNA Data Bank of Japan (DDBJ) under accession number DRA006953). piRNA-sized RNA species, approximately 30 nt long, were widely expressed in all tissues, and were used for ping-pong amplification analysis in *P. fucata*. Small RNA reads were pooled from similar tissues to perform piRNA annotation by unitas (v1.5.3), which was run with the option –pp^63^. In order to compare ping-pong signatures and the number of sequence reads within different tissue types, PPmeter (v0.4) was used to quantify and compare the extent of ongoing ping-pong amplification^26^. Pseudo-replicates were generated by repeated bootstrapping (default = 100) of a fixed number of sequence reads (default = 1,000,000) from a set of original small RNA sequence datasets. The ping-pong signature of each pseudo-replicate was then calculated, and the number of sequence reads that participated in the ping-pong amplification loop was counted. The resulting parameter (ping-pong reads per million bootstrapped reads, ppr-mbr) were subsequently used for quantification and direct comparison of ping-pong activity in different tissue datasets. Both tools were operated using default parameter settings.

### piRNA0001 silencing in *P. fucata* and quantification of piRNA expression

A single piRNA with the sequence 5’-UACUUUAACAUGGCACAGAUAUAAUGACCU-3’ (piRNA0001) presented the highest expression in *P. fucata* somatic tissues. To explore its function, an LNA-antagonist was used to silence its expression in *P. fucata*. Eight *P. fucata* oysters (approximately three years old) were randomly divided into two groups: LNA-antagonist group (LNA) and control group (Con). Oysters from the LNA group were injected with 100 μL 5mg mL^−1^ LNA-antagonist probe solution into their body cavity, while those in the Con group were injected with isometric PBS solution. The sequence of the high-affinity LNA-antagonist was 5’-AggTcaTtaTatCtgTgcCa-3’ (LNA in capitals)^64^. Two weeks after injection, all individuals were subjected to tissue collection. Somatic tissues, including from adductor muscle, gill, mantle, and gonad were collected and stored separately in 2-mL tubes containing RNA later (QIAGEN) overnight at 4 °C, and preserved at −80 °C until further use. Stem-loop RT-PCR was used for piRNA0001 relative expression analysis using *U6* sRNA as reference genes. This process was performed as described in our previous study^65^.

### Transcriptomic analysis of gene expression profiles after piRNA0001 silencing in *P. fucata*

Twenty-four libraries of somatic tissues, including adductor muscle, gill, and mantle, with four replicates for each tissue type from the LNA and Con groups, were constructed for RNA sequencing in 2017. Library construction, sequencing, *de novo* assembly, and functional annotation were performed as described above. Mapped fragments were normalized for RNA length according to transcripts per million reads (TPM)^66^, which facilitated the comparison of transcript levels between libraries. EdgeR was used to identify and draw the significant differentially expressed genes between the Con and LNA groups, at the following threshold: *P*-value < 0.05, and folds > 2^67^.

### Prediction of piRNA0001 targeting sites in *P. fucata*

piRNAs possess a second to eighth nucleotide seed sequence that requires near perfect complementarity between the piRNA and target genes, such as miRNAs^54^. Additional base-pairing outside of the seed is also important for piRNA targeting. piRNAs can only tolerate a few mismatches outside of the seed region^20,68^. In the present study, 3’ UTR sequences of differentially expressed genes were used for piRNA target site predictions. Because no piRNA target rules were explored in mollusks, we used miRNA-like pairing rules to predict potential target sites^53^. These were predicted between piRNA and mRNA 3’ UTR using miRanda tools (http://www.microrna.org/microrna/home.do)^69^ with an alignment score > 150 and energy < −10 kcal mol^−1^.

### Gene expression analysis by RT-PCR

To validate *PIWI* gene expression in *P. fucata*, the early development stage was achieved by artificial fertilization, and somatic and gonad tissues were dissected from one- and three-year-old oysters. Corresponding primers were designed based on the complete sequence of *P. fucata PIWI* genes (Table S4). β-actin was used as reference gene for *PIWI* relative expression calculation. First-stand cDNA was synthesized using the PrimerScript RT Master Mix reagent Kit (TaKaRa, Shiga, Japan), according to the manufacturer’s instructions. To determine the expression patterns of *PIWI* genes in various tissues at different developmental stages, all samples were analyzed using the Applied Biosystem 7300 Fast Real-Time PCR (RT-PCR) System (Life Technologies, California, USA). RT-PCR analysis was performed in a 96-well plate, where each well contained 20 μL of reaction mixture consisting of: 10 μL SYBR Premix Ex Taq II (TaKaRa), 0.4 μL ROX Reference Dye (50×) (TaKaRa), 0.8 μL of each primer (10 μM), 2 μL of complementary DNA (cDNA) template and 6 μL of sterilized double-distilled water. RT-PCR conditions were as follows: pre-denaturation at 95 °C for 30 s, followed by 40 cycles of amplification at 95 °C for 5 s and 60 °C for 31 s, a dissociation stage with one cycle at 95 °C for 15 s, and 60 °C for 60 s, and finally at 95 °C for 15 s after amplification. Each sample was tested in triplicate. The average value per gene was calculated from three replicates.

Nine differentially expressed genes, predicted to be targeted by piRNA0001, were also selected for relative expression analysis by RT-PCR as described above, with total RNA consistent with RNA_Seq2 libraries. Resulting PCR products predicted the interactio sites on mRNA 3’UTR. Relative expression of each mRNA quantified by RT-PCR was presented as n = 4, from three replicates. Ct values were represented by the mean values of three independent replicates, and the relative expression level of each gene was calculated using the 2^−ΔΔCt^ method^70^.

## Data availability

All sequencing data are deposited in the DNA Data Bank of Japan (DDBJ) database under accession DRA008674 and DRA007432.

## Abbreviations

piRNA: PIWI-interacting RNA
Ad: adductor muscle
Gi: gill
Ma: mantle
Go: gonad
*Piwil1*: PIWI-like gene 1
*Piwil2*: PIWI-like gene 2
Con: Control group
LNA: LNA-antagonist treatment group
RT-PCR: Realtime PCR
UTR: untranslated region
hpf: hours post fertilization
dpf: days post fertilization.

## Acknowledgements

We thank Yukihide Tomari and Natsuko Izumi for technical support of piRNA analysis. This work was supported by Japan Society for the Promotion of Science [Project number JP24248034] and Research Fellowship of Japan Society for the Promotion of Science for Young Scientists [Project number 18J13176].

## Author contributions

A.S., W.B. and K.S. designed the study. O.F. M.K., and N.K. provided the experimental animal samples. M and A.M. performed the transcriptomic libraries construction and sequencing with assistance from K.S. I. Yu. did the LNA-mediated piRNA silencing. H.S. and Y.K. performed all bioinformatic analyses, with support from I. Yo. H.S. collected and analyzed the data. A.S. and H.S. wrote the manuscript.

## Additional information

### Supplementary material

The supplementary material for this article can be found online.

