## Supplementary material for "Potential silencing of gene expression by PIWI-interacting RNAs (piRNAs) in somatic tissues in mollusk": Fig. S1-S7; Table S1-S2

**Supplementary figures:**

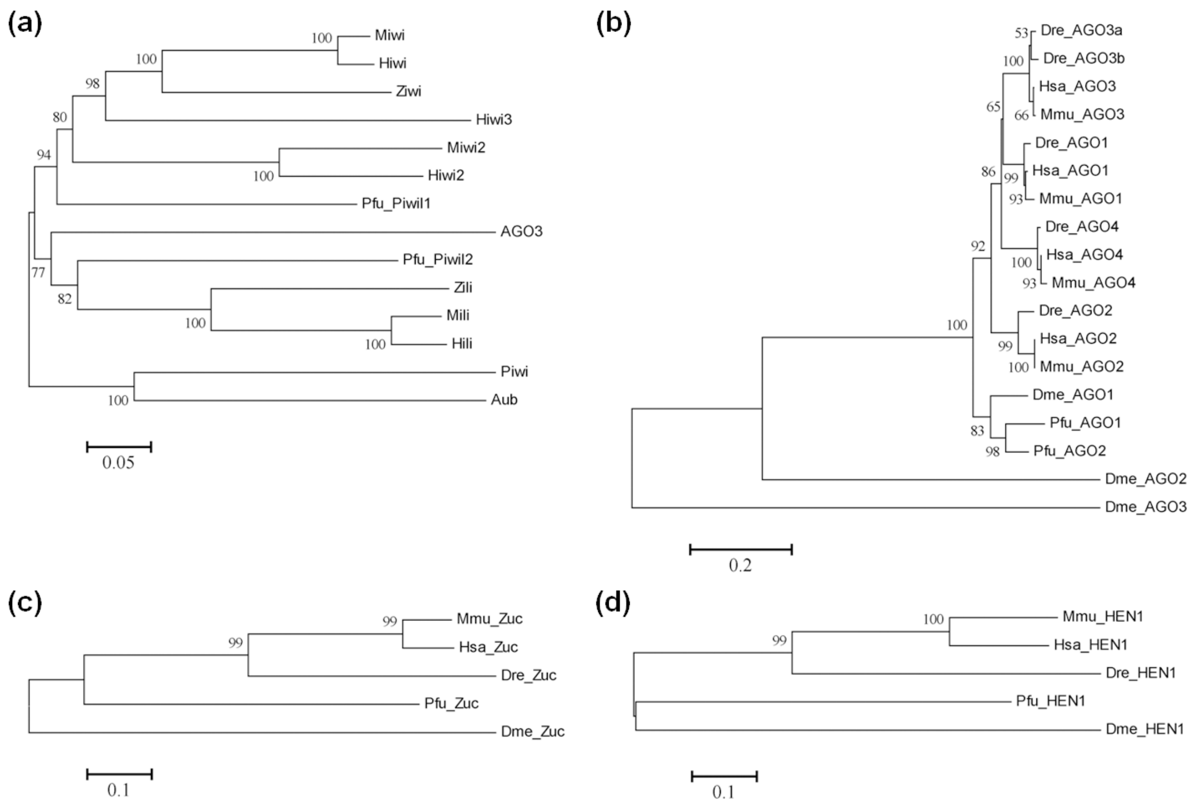


**Figure S1** Phylogenic tree of deduced amino acid sequences of piRNA biogenesis factors in *P. fucata* by neighbor joining (NJ) method using MEGA 5.0. The detail information of piRNA biogenesis factors homologues were listed in supplementary Table S5.

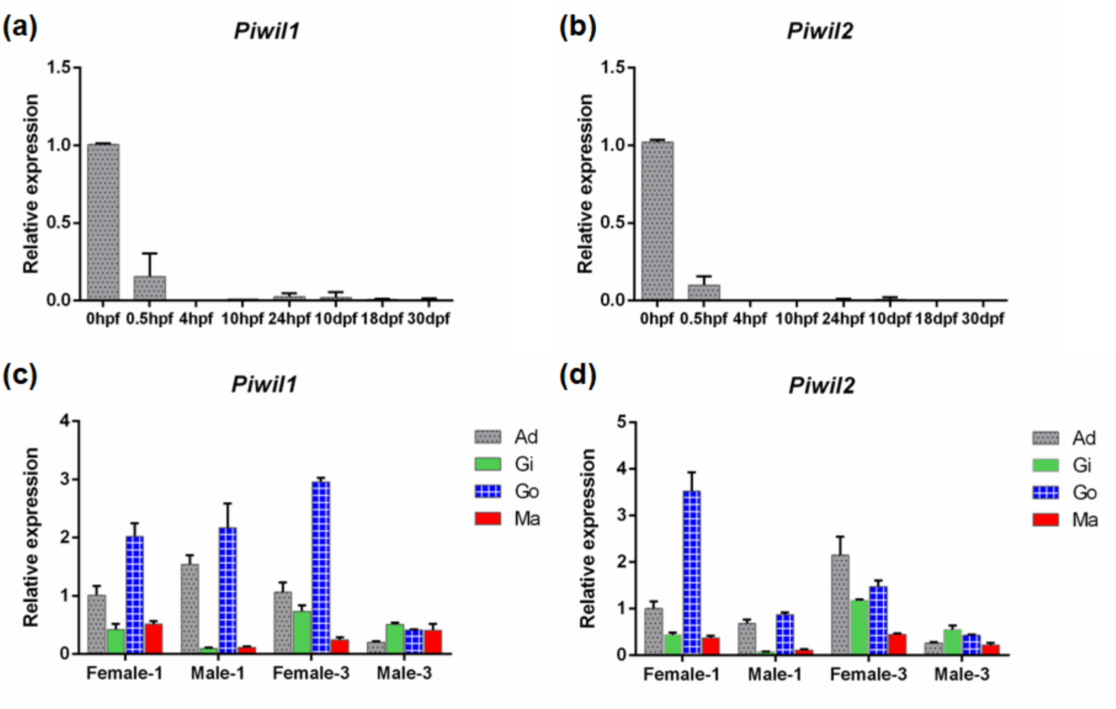


**Figure S2** Expression abundance of *Piwil1* and *Piwil2* at different developmental stages and tissues of *P. fucata*. *β-actin* was used as the reference gene. Developmental stages: 0 hour post-fertilization (hpf) (fertilized egg), 0.5 hpf (embryo), 4 hpf (blastula), 10 hpf (trochophore), 24 phf (D-shape larvae), 10 dpf (umbo larvae), 18 day post-fertilization (dpf) (pediveliger) and 30 dpf (spat). Tissues: adductor muscle (Ad), gill (Gi), gonad (Go) and mantle (Ma). The relative expressions at different developmental stages were normalized to the fertilized egg stage, while relative expressions in tissues were normalized to the expressions in adductor muscle at one-year old females.

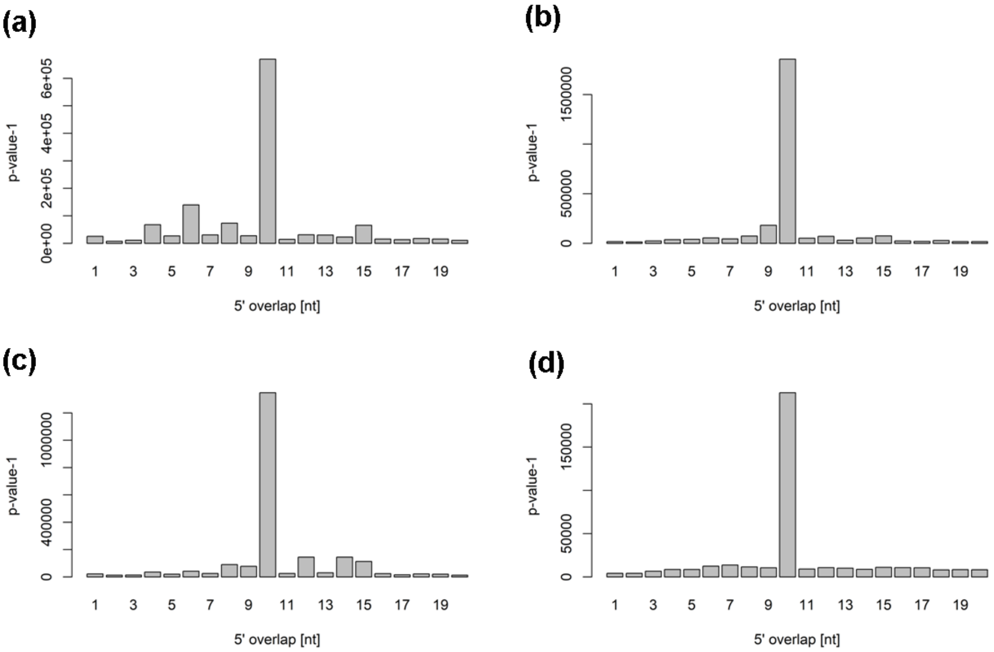


**Figure S3** Ping-pong signature of piRNA from *P. fucata* mantle tissue (a), adductor muscle (b), gill (c), and gonad tissues (d).

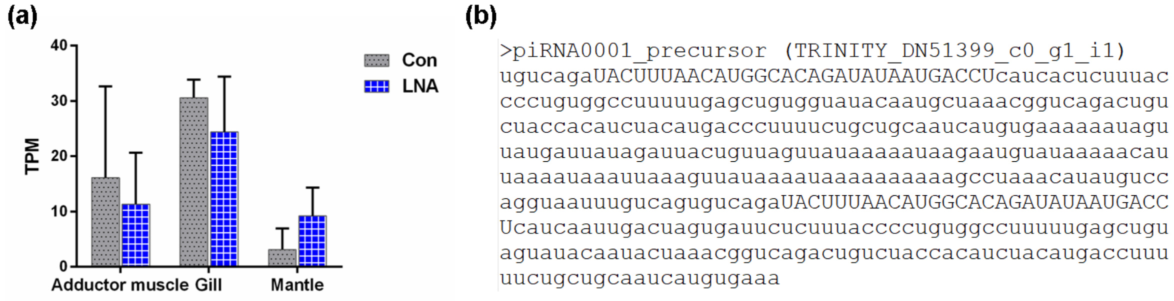


**Figure S4** Expression profiles of piRNA0001 precursor and piRNA0001 loci on precursor. (a) Expression patterns of piRNA0001 precursor by transcriptomic analysis. TPM: transcripts per million reads; (b) Sequence of piRNA0001 precursor and piRNA0001 loci (in capitals).

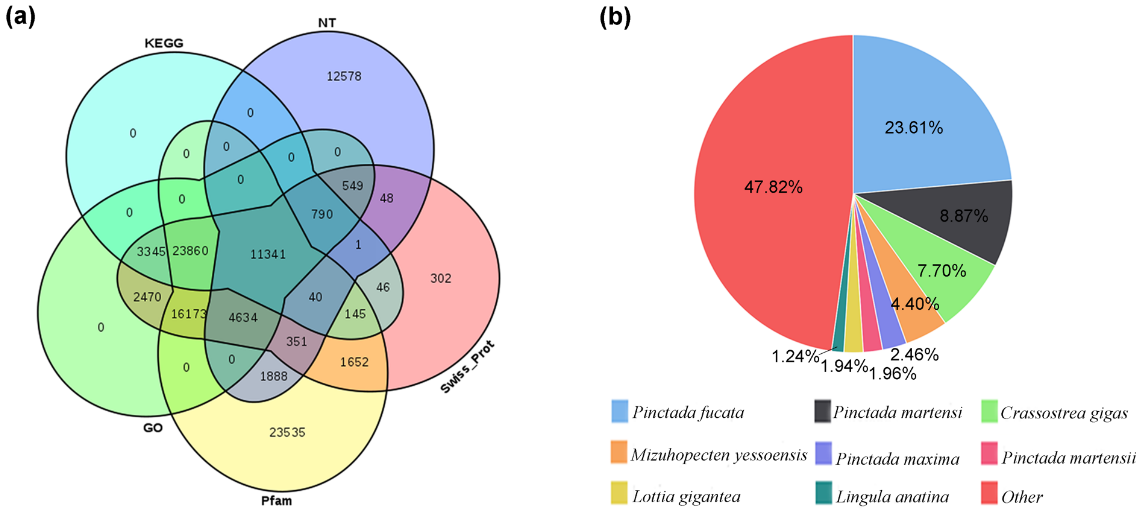


**Figure S5** Annotation results of transcripts in *P. fucata*. (a) Detection of homologous genes in five public databases (NT, Swiss-Prot, Pfam, GO and KEGG). (b) Proportion of species annotated for these transcripts in *P. fucata*.

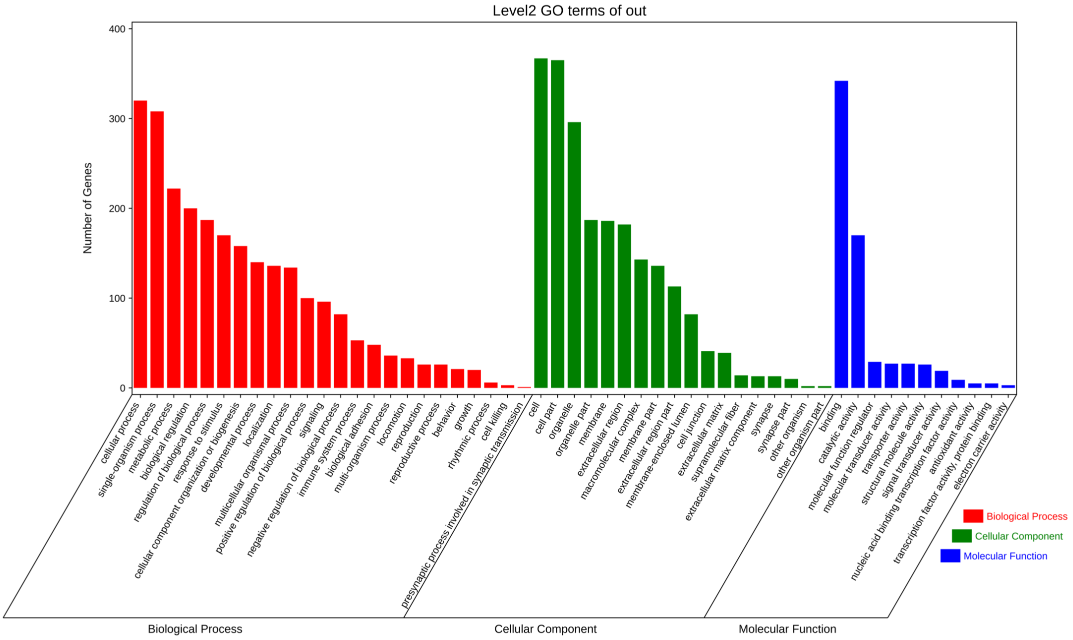


**Figure S6** GO enrichment analyses of differentially expressed genes after piRNA0001 silencing in the pearl oyster *P. fucata*.

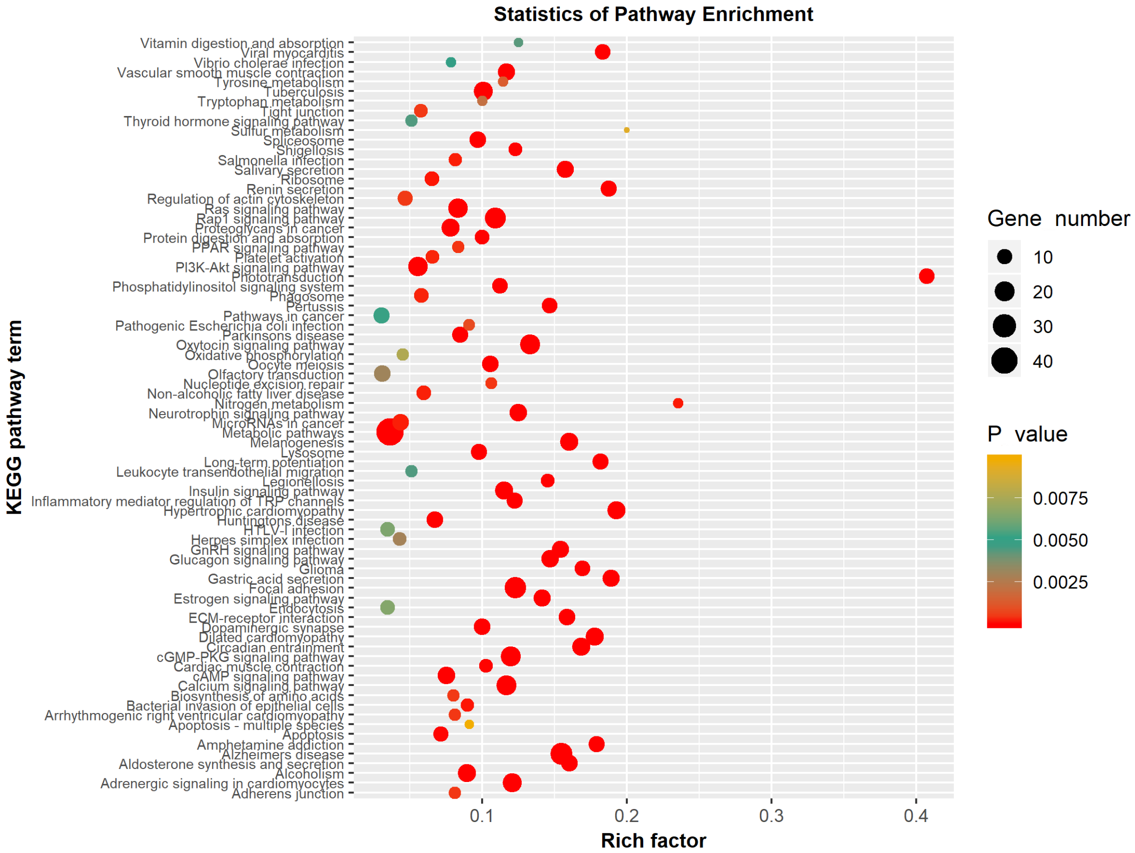


**Figure S7** KEGG enrichment analyses of differentially expressed genes after piRNA0001 silencing in the pearl oyster *P. fucata*. Rich factor is the ratio of the number of DEGs to the total number of genes in a certain pathway. The size and color of the dots represent the gene numbers and p value, respectively.

**Supplementary Tables：**

**Table S1** Statistics of sequencing data of transcriptomic analysis of piRNA biogenesis factors in *P. fucata*.

| Library | Clean Reads | Clean bases | Q30 | GC content |
|---|---|---|---|---|
| F1Ad | 12138234 | 1820735100 | 96.90% | 43.00% |
| F1Gi | 17637746 | 2645661900 | 96.64% | 41.00% |
| F1Go | 6219058 | 932858700 | 96.51% | 43.00% |
| F1Ma | 14067508 | 2110126200 | 95.99% | 43.00% |
| M1Ad | 12433782 | 1865067300 | 96.95% | 43.00% |
| M1Gi | 14214052 | 2132107800 | 96.45% | 41.00% |
| M1Go | 14879724 | 2231958600 | 96.70% | 41.00% |
| M1Ma | 9909510 | 1486426500 | 96.20% | 43.00% |
| F3Ad | 15919858 | 2387978700 | 96.95% | 43.00% |
| F3Gi | 13790444 | 2068566600 | 96.65% | 40.00% |
| F3Go | 5327732 | 799159800 | 96.56% | 42.00% |
| F3Ma | 9746006 | 1461900900 | 96.33% | 43.00% |
| M3Ad | 5876794 | 881519100 | 96.85% | 43.00% |
| M3Gi | 13176792 | 1976518800 | 96.88% | 41.00% |
| M3Go | 15106294 | 2265944100 | 96.78% | 40.00% |
| M3Ma | 10668480 | 1600272000 | 96.46% | 43.00% |
| Total | 191112014 | 28666802100 | 96.61% | 42.06% |

**Table S2** Statistics of sequencing data of transcriptomic analysis of gene expression profiles after piRNA0001 silencing in *P. fucata*.

| Library Name | Clean Reads | Clean bases | Q20 | GC content |
|---|---|---|---|---|
| Con_Ad1 | 4887098 | 488709800 | 98.28% | 43.36% |
| Con_Ad2 | 7202438 | 720243800 | 98.16% | 43.00% |
| Con_Ad3 | 16224294 | 1622429400 | 98.12% | 42.12% |
| Con_Ad4 | 6528166 | 652816600 | 98.26% | 42.14% |
| Con_Gi1 | 8917184 | 891718400 | 98.11% | 40.92% |
| Con_Gi2 | 5464414 | 546441400 | 98.20% | 41.38% |
| Con_Gi3 | 10260062 | 1026006200 | 98.24% | 39.70% |
| Con_Gi4 | 6645710 | 664571000 | 98.25% | 41.23% |
| Con_Ma1 | 9400386 | 940038600 | 98.40% | 44.84% |
| Con_Ma2 | 16254080 | 1625408000 | 98.14% | 43.25% |
| Con_Ma3 | 12874756 | 1287475600 | 98.05% | 43.40% |
| Con_Ma4 | 4499214 | 449921400 | 98.29% | 42.91% |
| LNA_Ad1 | 17189860 | 1718986000 | 98.11% | 43.21% |
| LNA_Ad2 | 3305308 | 330530800 | 98.28% | 43.36% |
| LNA_Ad3 | 12267128 | 1226712800 | 98.29% | 43.53% |
| LNA_Ad4 | 9161644 | 916164400 | 98.26% | 43.26% |
| LNA_Gi1 | 16616448 | 1661644800 | 98.23% | 42.85% |
| LNA_Gi2 | 14301690 | 1430169000 | 98.16% | 40.96% |
| LNA_Gi3 | 11188624 | 1118862400 | 98.29% | 39.45% |
| LNA_Gi4 | 12911558 | 1291155800 | 98.16% | 42.90% |
| LNA_Ma1 | 23730166 | 2373016600 | 98.23% | 43.75% |
| LNA_Ma2 | 15261694 | 1526169400 | 98.39% | 43.07% |
| LNA_Ma3 | 11829700 | 1182970000 | 98.12% | 42.86% |
| LNA_Ma4 | 8784180 | 878418000 | 98.20% | 42.93% |
| Total | 265705802 | 26570280200 | 98.22% | 42.52% |

**Table S3**Differentially expressed genes between Con and LNA group in *P. fucata*

**Table S4** Primers used in the present study

| Primer name | Primer sequence (5' to 3') | Usage |
|---|---|---|
| Piwil1_F | TTACAAGACGCAACAGTGCTCAG | RT_PCR primer for *Piwil1* |
| Piwil1_R | GCCTTTAGATAAGTGTCATTTCGG | |
| Piwil2_F | AGGGCTTACTGATGAACTGAGGA | RT_PCR primer for *Piwil2* |
| Piwil2_R | CCCATTTTGTCAACTCTGCTTTC | |
| β-actin_F | CTGTTCTGTCTTTGTATGCTTCC | RT_PCR primer for *β-actin* |
| β-actin_R | TCAGGTAGTCGGTGATGTCTCT | |
| piRNA0001_SL | CTCAACTGGTGTCGTGGAGTCGGCAATTCAGTTGAGAGGTCATT | Stem-loop for piRNA0001 |
| piRNA0001_F | ACACTCCAGCTGGGTACTTTAACATGGCACAGATATAA | RT_PCR primer for piRNA0001 |
| piRNA0001_R | TGGTGTCGTGGAGTCG | |
| U6_F | TTGCTTCGGCGGTACATATA | RT_PCR primer for *U6* |
| U6_R | ATTTGCGTGTCATCCTTGC | RT_PCR & stem-loop primer for *U6* |
| ART2_F | ATTGGAGGGCAAGTCTGG | RT_PCR primer for *ART2* |
| ART2_R | CGGTCAGGTAGGTCAGGAT | |
| CBF_F | CTAATGGAGTTATTGGCTG | RT_PCR primer for *CBF* |
| CBF_R | TAGGTTGTCTAAGCACTGA | |
| ESI1L_F | ACAAGCAATGTGCGTATGA | RT_PCR primer for *ESI1L* |
| ESI1L_R | ACCATTTGGGCATAGTGTC | |
| NBR1_F | GTCCATTTATGCTTATCTCAC | RT_PCR primer for *NBR1* |
| NBR1_R | AACTTTTTTCTAACAGCCAT | |
| PA2HB_F | AGATTTGAGGGTGAAGATTAT | RT_PCR primer for *PA2HB* |
| PA2HB_R | AAAGATTTTCGTTATTGTTGA | |
| SRS10_F | TTATTACTCACGAAATCCCAG | RT_PCR primer for *SRS10* |
| SRS10_R | TTAACTTTTCCTTGTGAGCTT | |
| TYR1_F | ATGTCGTTTTGCTCTAAGTC | RT_PCR primer for *TYR1* |
| TYR1_R | GAGCGAATCTTTGGTAGTAT | |
| VINC_F | CAGTATGGTGAACAAAGGAAGTC | RT_PCR primer for *VINC* |
| VINC_R | TTTGTCTTGCCTTTTTTTCCT | |
| ZNF622_F | ATGCCTGGATTATTTATTTTG | RT_PCR primer for *ZNF622* |
| ZNF622_R | TGGGTGTCTGAATACGAAAAC | |

**Table S5** GeneBank accession numbers of PIWI protein sequences used in phylogenetic analyses

| Species | Protein symbol | NCBI Accession |
|---|---|---|
| *Homo sapiens* | Hiwi | AAC97371.2 |
| *Homo sapiens* | Hili | NP_001129193.1 |
| *Homo sapiens* | Hiwi2 | NP_689644.2 |
| *Homo sapiens* | Hiwi3 | NP_001008496.2 |
| *Mus musculus* | Miwi | BAA93705.1 |
| *Mus musculus* | Mili | BAA93706.1 |
| *Mus musculus* | Miwi2 | NP_808573.2 |
| *Danio rerio* | Ziwi | NP_899181.1 |
| *Danio rerio* | Zili | ACH96370.1 |
| *Drosophila melanogaster* | Piwi | AGL81535.1 |
| *Drosophila melanogaster* | Aub | NP_476734.1 |
| *Homo sapiens* | Has_AGO3 | NP_079128.2 |
| *Homo sapiens* | Has_AGO4 | XP_005270636.1 |
| *Mus musculus* | Mmu_AGO1 | XP_011238823.1 |
| *Mus musculus* | Mmu_AGO2 | NP_694818.3 |
| *Mus musculus* | Mmu_AGO3 | NP_700451.2 |
| *Mus musculus* | Mmu_AGO4 | NP_694817.2 |
| *Danio rerio* | Dre_AGO1 | AFU66007.1 |
| *Danio rerio* | Dre_AGO2 | AFU66008.1 |
| *Danio rerio* | Dre_AGO3a | AFU66009.1 |
| *Danio rerio* | Dre_AGO3b | AFU66010.1 |
| *Danio rerio* | Dre_AGO4 | AFU66011.1 |
| *Drosophila melanogaster* | Dme_AGO1 | NP_725341.1 |
| *Drosophila melanogaster* | Dme_AGO2 | NP_001261882.1 |
| *Drosophila melanogaster* | Dme_AGO3 | NP_001036629.2 |
| *Homo sapiens* | Has_AGO1 | NP_001304052.1 |
| *Homo sapiens* | Has_AGO2 | NP_036286.2 |
| *Homo sapiens* | Has_Zuc | NP_849158.2 |
| *Mus musculus* | Mmu_Zuc | NP_001277212.1 |
| *Danio rerio* | Dre_Zuc | NP_001082883.1 |
| *Drosophila melanogaster* | Dme_Zuc | AAF53139.1 |
| *Homo sapiens* | Has_HEN1 | NP_001096062.1 |
| *Mus musculus* | Mmu_HEN1 | Q8CAE2.1 |
| *Danio rerio* | Dre_HEN1 | Q568P9.2 |
| *Drosophila melanogaster* | Dme_HEN1 | NP_610732.1 |

**Supplementary sequences:**

The front and rear lowercase letters represent the 5'UTR and 3'UTR of the genes, respectively. The uppercase letters represent the open reading frame including the stop codon. The red and yellow frame shows the predicted piRNA:mRNA interaction sites and the polyA signal on 3'UTR, respectively.

>TRINITY_DN58940_c1_g1_i1 ART2
gcgagtcggcttgcggttttccgacttcgtcgggcgtttcttataaagctgctgcctgcgtcATGGAACGTCCGATCTCCGGTCCCACGCCGGCTATGCCGCGCAAATA
TGATTTGCGTAGGAACTGTGGCACAAAATCCGCATGGGGAAGATGTTCATTCCCCAAAATTTTCAGGCAAACTAGTGGCTGCCC
GGCCCGAGGCTTTCGAGTTTCCGGCATGCAGTCCGATACGGTTTCGGTCCCGAGGAACGACGTCGGTCTACAGAGCTATCCGCT
CTTAACGACCCGGCTGTTCCGAGGTGGTCGCCTCTCGTCGAGATGTGCGATCCGCGTGTAGaaacagggcacggtgtccgttactcccatatctggag
ggagcgaatcggaggcctgccttgcgtctaagagccagctcggcactccgccgagcacacggccctggacgacttcgattgagcccgtggacggcacggcgatggctgggaagttctgtatcaccccggc
ggacgctgtaacgcctcctcggctgctgtaatcgcataaccgatcccggtacagtccttgtttgactgaactcagggaacggggacggggcggcttcggccgctgtcgtcgaatagctatctggttgatcctgc
cagtagtcatatgcttgtctcaaagattaagccatgcatgtctaagtacagacttatacatagtgaaaccgcgaatggctcattaaatcagttatggttccttagatcctacaatcctacttggataactgtggcaattct
agagctaatacatggcaaaacagctccgacctcacgggaagagcgcttttattagaacaaaactaatggcggtcgcaaggccgtctcaccttggtgattctgaataactttgtgctgatcgcatgagcttgcctc
ggcgacgtatctttcaaatgtctgccctatcaactttcgatggtacgtgctatgcctaccatggttgtaacgggtaacggggaatcagggttcgattccggagagggagcatgagaaacggctaccacatccaa
ggaaggcagcaggcgcgcaaattacccactcctggcacggggaggtagtgacgaaaaataacaatacgggactcttacgaggccccgtaattggaatgagtacactttaaaccttttaacgagtatcaattgg
agggcaagtctggtgccagcagccgCGGTAATTCCAGCTCCAATAGCGTATATTAAAGTTgttgcgtttaaaaagctcgtagttggatctcgggtccaggcttgcggtccgctc
cttggcggtcactgcttgatcctgacctacctgaccggtttttgcccttggtgctcttgattgagtgtctcgggtgaccggaacgtttactttgaaaaaattagagtgctcaaagcaggcgatttgcctgaataatggt
gcatggaataatggaagaggacctcggttctattttgttggttttcggaaccagaggtaatgattaagagggactgacgggggcattggtattgcggtgttagaggtgaaattcttggatcgccgcaagactaac
agctgcgaaagcatttgccaagaatgttttcattaatcaagaacgatagtcagaggttcgaagacgatcagataccgtcgtagttctgactttaaacgatgccaactggcgatccgccggagttgcttcaatgact
cggtgggcagcctccgggaaaccaaagtttttgggttccgggggaagtatggttgcaaagctgaaacttaaaggaattgacggaagggcaccaccaggagtggagcctgcggcttaatttgactcaacacg
ggaaaactcacccggcccggacactgtaaggattgacagattgagagctctttcttgattcggtgggtggtggtgcatggccgttcttagttggtggagcgatttgtctggttaattccgataacgaacgagactc
tagcctactaaatagttcgccgatcacaaatgctgtcggcgccaacttcttagagggacaagtggcgtttagccacacgagattgagcaataacaggtctgtgatgcccttagatgttcggggccgcacgtgcg
ctacactgaaggcatcaacgtgtctgtcccctggcccgagagggttgggtaacccgttgaaccgccttcgtgctagggattggggcttgtaattattccccatgaacgaggaattcccagtaagcgcgagtcat
aagctcgcgttgattacgtccctgccctttgtacacaccgcccgtcgctactaccgattggttggcattgtgagcccctcggattggtcccggcagcggggtttctcgcagccggtgtgccgagaagaagggcg
aacagttctggctagaggaagtaaaagtcgtaacaaggtttccgtaggtgaacctgcggaaggatcattaacaaaaactcagcaagagtgaaaacttgcgctgaatgagatcattacgaactcaaagcggtgg
gagtgaaggacgtaacgtccgaggtactcccctgcgaacctatacgatgtgctcgttgGGGTACCCAGATCTGGAGGGTTTAAAGTGtggtctgtatgtaaccgccccgtttcaaca
agacgccagtagtgggtacttgggccggtctgaatctgtgttgcagatcggcgagagtccgcttctccgaagtggttaggtttaaagagcggcccgtttctggtcagggccgccgcccggcgtctctcaaacct
ttttttcgtactgttgatacaaactagcggtgacttcggtcgccgggtcgcatcctaacagatcgaccgaatttcataaagcaaatacaactctaagcggtggatcactcggctcgtgcgtcgatgaagagcgcag
ctagctgcgtgaattaatgtgaattgcaggacacattgaacatcgacatcttgaacgcatattgcggcctagggtcactcccttggccacgcctgtctcagggtcggtgaaacatcttcgcaagaaattgctctgta
tgcttgcgcattgggctgtcgaagtacttcggtactccgtcgcctcaagttcagacccgattgcctcggcacgtcgtctgacggtctcgaccattcatgacgtagctcgctgttctgtttgaagcactgcgccggtc
caagactgtcgatcgatgaccacgggagcgatcgaaactattcatttaggtttcacacctctctcatccgacctgagatcagacgagacaaccccctgaatttaagcatatcactaaggggaggaaaagaaact
aactaggattccctcagtaacggcgagtgaagcgggaaaagcccagcaccgaatccctcggcctcgcgccgctgggacctgtggtgtttaggacgcctcactgcggtggggtacgggtgcccaagtccttc
tgatcggggcctctcccatagcgggtgtcaggcctttacgggcactcgtcccttctgctaagaacgtccttggagtcgggttgtttgagaatgcagcccaaagtgggtggtaaactccatctaaggctaaatact
ggcacgagtccgatagcggacaagtaccgtgagggaaagttgaaaagaactttgaagagagagttcaagagtacgtgaaaccgcttagaggtaaacgggtggacccgcaaagtcggcccggggaattca
actctctgcccagtcggcgggcggtgcgcttgggatccgtgaggactcggcgtgctgctaaggtcggcggcggggtgtgcactttctccgggtagagcgccacgaccggtttgctggcggcgagaaaccg
gagaggaaggtgacctgcttgcttcggcagtaggtgttaagactctccgcagttcgctcgtcggcggaccgaggatctgccgcggttcatctgcccctggttctccggatgcgatcgaccggccttcactgctt
gcagtgctggccaaccgcgtgtctggactcaggccgggaggcacggaccgctttcgggtctgtggcgaatcggtcggtactccacccgacccgtcttgtaacacggaccaaggagtctaacatgtgcgcga
gtcatggggttttacgaaacctaaaggcgcaatgaaagtgaaggcagcctcgggtttgcctaggtgggatcctctccctcgcggggaggcgcaccaccggcccgtctcgtccgcattgtcggtgaggcgga
gcaagagcgtacacgttgggacccgaaagatggtgaactatgcttgagtaggacgaagccagaggaaactctggtggaggctcaaagcggtactgacgtgcaattcgttcgtggaacttgagtataggggc
gaaaaaccaatcgaaccatctagtagctggttccctctgaagtttccctcaggatagctggcatccagggcagtttcatccggtaaagcgaatgattagaggccttggggacgaaacgacctcaacctattctca
aactttaaatgggtgagaagttagactcgcttaattggagtctagccttcgaatgcgtggtgccaagtgggccacttttggtaagcagaactggcgctgtgggatgaaccaaacgcccggttaaggtgcctaac
gctgacgctcatcagataccataaaaggtgttggttgatacagacagcaggacggtggccatggaagtcggaatccgctaaggagtgtgtaacaactcacctgccgaatcaactagccctgaaaatggatgg
cgctagagcgtcggacctataccgggccgtcgccgcaatacgagcaccccgggtgccaagcggcgacgagtaagagggccgccgcggtgtgcactgaagcctggggcgtgagcctgggtggagccgc
cgcgggtgcagatcttggtggtagtagcaaatattcaaacgagaactttgaagactgaagtggagaagggttccatgtgaacagcagttgaacgtgggtcagtcggccctaagagataggaaaactccgttct
gaaacggggcactgtctcaatctgaaagtacatgttgcctgtcttctatcgaaagggaatcgggtcaatattcccgaacctggcatcggagaatgctcctcacggggcagtgtgcggtaacgcaaacgaactc
ggagacgtcggcggaagtcccggtaagagttctcttttcttattaagggacggactccctggaatcggcttggccggagatagggacgttgttcccgtaaagcaccgcctctcttgcgtgtgtccggtgcacttc
cgacgacccttgaaaatccgagggagacaactctcgtgccagatcgtacccatatccgcagcaggtctccaaggtgcacagcctctagtcgatagaacaatgtaggtaagggaagtcggcaaattagatccg
taacttcgggaaaaggattggctctaagggctgggccagtcgggctggagtacgaagcaggatcgggacgggcacgggctgggcgagaccttcgggtcgagctcggaccgctgtctcaaccgtcttgtgg
actgcctcagctatgtcgtggctcctcgctgggtcgcggcttcgtctggcaactaacagccaacttagaactggcacggaccaggggaatccgactgtctaattaaaacaaagcattgcgatggccgtcaccc
ggtgttgacgcaatgtgatttctgcccagtgctctgaatgtcaaagtgaagaaattcaatcaagcgcgggtaaacggcgggagtaactatgactctcttaaggtagccaaatgcctcgtcatctaattagtgacgc
gcatgaatggattaacgagattcccactgtccctatctactatctagcgaaaccacagccaagggaacgggcttggcagaaacagcggggaaagaagaccctgttgagcttgactctagtccgactttgtgaa
gagacatgattggtgtagcataggtgggagcttcggcacaattgaaataccactactttcatcgtttctttacttattcagtaaagcggagagcggggcgcaagcctctcgattctggcattaagcccccggccttc
gttgtcgggggtgatccgctctgatgacagtgtcaggcggggagtttgactggggcggtacatctgtcaaacggtaacgcaggtgtcctaaggtgagctcagcgaggacggaaacctcgcgtagagtaaaa
gggcaaaagctcacttgattttgattttcagtacgaatacagaccgtgaaagcgtggcctatcgatccttttgattttaagagttttaagcaagaggtgtcagaaaagttaccacagggataactggcttgtggcag

ccaagcgttcatagcgacgttgctttttgatccttcgatgtcggctcttcctatcattgtgaagcagaattcaccaagcgttggcttgttcacccactaatagggaacgtgagctgggtttagaccgtcgtgagacag
gttagttttaccctactgttgactagcgtcgttgcaatggtaatcctgctcagtacgagaggaaccgcaggttcagacatttggtttatgtacttggctgataagccaatggtgcgaggctaccatctgagggattat
gactgaacgcctctaagtcagaatctcgcccaaatttacaacgatactttctatgcgctccaggccctgggaggcaacgataaacggggtaccgtcccggcaacgggcgtccccggtggttaagccacagta
caggaccacatgcaggcactgactttcgtgtctgtgggtctttacgatatccatcccatgcggtgtcggagcgtaaaatcattcgtagacgacctagttcttggtcggggtgtcgtacctagtagagcagccacct
cactgcgatctattgagactaagcctttcgactagccgatttgtccgcttaagacggactatgtcaattaagctattttcgaaagtttgatttgtgacgagttgtgtgtacttttatcctggtggctaatcatacatggtttc
aaaaattcattagatgtatacttagatgaatgaatatgtaacttttaacatgtgtgaacttgtgaagtactgtgtttatttgattaattggagtataaatctgactaaaacatattatgtacagaacacac
>TRINITY_DN46965_c0_g2_i2 CAH14
cttagccaaccaaatagcagttcaaatctgtaattggcacttgatcgatatccaccaatcagaatgtagtattacaaccattccttcctccgtccaacattatctgataacaggttttttttctgaggctgtcatccacgct
ttgagctctgaccccaacactactataccATGGAACTGACAGTACCTCTGATTCTTTGTGTGACAGGGATCTTATCTCTTACTAAAGCTGCAGAAT
GGGGTTACACAGGATCAAATGGACTTCATCATTGGCCGTCGTCTTTCCCAAGTTGTGGTAAACACAGCCAGTCTCCCATTGATTT
CCCAGAGGAGAGTGAAATGGAACTTGACGTCAACCTGAAACCAATGAAAATTACAGGACTTGAGAGGACCAAAAACTACAGA
CTGGAGCTCTTTAATAACGGTCATTCAGCTCAAGTGAATGTAGATGGTGACGTCTATGTAGAAGGAGGGGGACTGCCAAATAGA
TTCCGAACTGCTCAGTTTCATTTCCACTGGGGACGTGATGACAACACTGGTTCAGAACATCTCTACAGCGGGGGCTCATTCCCA
TTAGAGCTCCACGTAGTAAACTACAATGTTAAATATGGAAACAGTTCAAATGCCATGACAAAAGTAGGCGATGGATTGGCTGTC
CTAGGATTCTGGTTCGAGCGTTCTGATTCTGACAATCCTGTTCTGACTCCGCTTATCAACACACTTTCTTCTGTCCAAAATGCAG
ATCCTTCAACAAAGGTTCCCGTTAGTGGTCTGGACCTTGGTAAGGTCATATCAGGGAACTTGGATCGCTTCTACAGATACTCCGG
TAGTCTCACAACGCCACCATGCTACGAAAGTGTCATCTGGTCGGTATTCCAAACAAAAATACCAATCTCATCAAGTCAGCTTAC
AGAATTTAGAAGACTGAAAAAGGCTCCACATGGAGGGAAGACCACAGCTCTGGTGGATAACTACCGGCCTCCGAAAACCATC
AACGGAAGAAAAATCTCCATCAGCTTCCCGTCTGGTGTCGCCACTGTCTCTGCGAGTAGTCACGTGATATCAATATTATTTTGTA
TCTTTTTATTAGCAATAAAGATGACATAGgcttgtgaaatcacaaaaatcacaaagacagaattccgattttataataatgtgtataccataatcagaaatcatacagtgccaattacttga
aaatctgccgttatattgtgccaattacaatatagtcccaacttttactgttgatgacttgttgaaaaccacaaatataatattttacaataaaactattatacaaaggatttgaaacaaaaatgtgccatattcgcaaaac
tagtcatgctgtaatcatgctttggtgttagcttcgataatgttctatgtatgtagcaatgtataATTTGATGTATGTACATGTACTTATTTTTAAAGTAtttgcaaatattttttcaagaact
gcgttcattctgaatcaaatgtgttcacaatcacttacgtatgctctctaaactgtgatttacaatatcatataatttcatataaattcataaaatgtgttatctattaaat
>TRINITY_DN36651_c0_g1_i5 CBF
gttgcggtttagtaatgaatgaaggggtttgagaaagcATGCTATCACTTCTTACGAACACGTTGTGTTGTAAAGTTGGAAATAATTTCGCCACAAACA
TGCCCCGAGTGGTGCCGGATCAAAGATCGACTTTTGAAAACGACGAACTTTTCAGGAAATTAAGCAGAGAATGTGAGGTAAAG
TACACTGGTTTTAGGGATAGACCCACAGAGGAAAGGCAGCTGAGATTTCAAACAGAATGTAGGGAGGGTCACGCTGATGTGGC
TTTCGTAGCAACAGGAACCAATTTAAATTTATTATTTGAATCAAATGCATGGTCAGAGAGGGATGAAGATAGAGTGGTGACCAG
GGAATATGTAGACTTTGACAgggaacctggaaaggtccatttaaagtcCCGTTTTATAATGAATGGAGTTTGTGTGGTGTGGAGAGGCTGGTTGG
ATTTACAAAGGTTAGATGGGGTAGGCTGTATGGAATTCGACTCAGAACGAGCCGAGGTTGAGGATGCCTTATATAGAGAACAGA
GAGATGAGTACAATCGGAGATTACGAGATTTTCCCGACATACAGCGGCAATATCGAGAGCAGCAAGAGCGTCGCGACCAGGAA
GCAGAGGTAGTGGAGGCTTTGTTGTGTATATCTGAGGATCGTAAATCGTAAccccaactttcgttctccatatttgtggttttgtttatttgaccttatttcatcccattct
gttgtgttccatgtgataacttactttaaaatgactcaaactgaagtgaaaaagagcttaaaagaaataccaagattacatacatagaccaagttaatgctggactgtcaaaatttcactttatttctctaactttctataa
atctgtcaatttcagggtggtttttaagaaactctaccataaactttcttgctcaaatctgattggacaagatttaaagcagccaactctattatccaatcagctgtgacatgacataaagtatatgacaaatcttctactt
accgtctcagcatttcatcctatgacaatttttgttattttgatgttaaagattatttacttttgcatgtatgtttgtgtatgcttaaagcattgtgataaaatgttatatataaggtattgttggtatcttaagtttcatcattatttc
atacatttatgcaataccattattttaTCATTATTACCATATGTTGTATTTTAAAGTGagtttctattgttgttttatagatattttaagatgttttataaaatgataaacttctgtttaaagagtgata
tgatttgtttctacttgcttaaaccaatcaacaggtgtgcttcaaatcctaatggagttattggctgatatttttcaaatcCTTGCAAGATGATTGGCTAATGTTAAAGTTtaaaattttattttg
cacttgcttgtgaaagctgaattgtagtggaactttaaagtcagtgcttagacaacctaattttgcttcttgtcataattatttttgtatgttttgatgttgtgttttcatagattatgaatgtgatattgtatgaattcaggaaaa
aaaggaactttccattgaattgaatatgccaatcattatttcatatatcattttaatttcatgtactattccatttttatatgagaaatgttttacattgacttcatagtgtaatcatgagaaactgttcagtcattatcatgaaat
gaccaatctagtcaattgcatatatattgatgtcacagtttcttgacttgctcatatgcaatgccaaatgcaatatgtacatagaattatctcaaaagtgtatgtcttttattacattatatagaatacagaggaaatatcag
atctgtttatatatatgatataattttagctcctttcaccctgtgatgaaatgaatttatattttgaaaataaagtatgaataag
>TRINITY_DN42088_c0_g1_i1 CDC42
cgaagtaccgcggtgtgcattcgacatattttcgtggttttaacgaagacacctcaatatgttaagtatgaaatggtacacttgaaactgtatcctattatcgaatgacatagtacatcaattgtaactttctttatgtatc
gtaaagtttaagggaggtcttaaccggaaatacggaagtgcctatcgtcagccgacgtcacaaaacaaagatgctacacacttgaccagcttgatgatgtagtatgtaaatgattaaaacttgtgaaattttactaa
tcggtcattgcgctatttaaaaTTTTAACGAAACTTCAAACATATTAAAGTGtgtggataaaatttttaagatttaaaagctacggaaactgaaagaaggaagtataattcaaacggaat
aacaaatctgaaaacagcgtggtgatagtaaattaaactagttttgaattcaaacatttttttggaatttgaaATGAAGATGGAAATTGAGTCTGATGAGACTGCGGATGGTG
CTAATGTACCCCAGGATAAAAATAAAAACGAACTGACCGATGTTTTAAAGGCACGGGCTGGAGAAGGGGAGATCAAATGTTGT
ATAGTCGGTGATGGCGGTGTGGGCAAGACGAGCATGTTGCTGAGCTACACGACCGGCAAAATGTTGACAAATTACGAACCAAC
ATGCTTTGAAAGTTATGTCATTGAGGTGAAACAGGGGGAAACCACGAAGAAGATGTTTGTAATGGACACAGCTGGTCAAGAAA
CCTACGACAGATTAAGGACACTTTCATATTACGACACAGACGTATTTATAGTGTGTTTTTCGGTTGACGATCAGGACTCATTTGA
CAATGTGAGAACGAGATGGATACCAGAAATCAAAGAATTCCGACCGAATACTCCCTTTATCATGGTAGGTACCCAAACAGATCT
CAGACACTCAGTTGTGGACATAACAGACGCTTGCGTATCATCAACGAGAGGCAAAAAAGCTGCAAAGAAACTAGGTGCCAAA
AAATATATCGAATGTAGTTCAGTAGACGGACAAGGTACGGACTATGTATTCGAAGTGGCACTAAAGACGGCGCAATCTCCCATG
GGACAGAAGAAAATATGGAAAAGCATCAAGAGAGCTTTCCGGAAAAAAGATGCTGCAGTTTGAtaactgcatctttctggttaattcgttatccgga
atttcctgctttcccacattcggattgaaagtgttatatcaccgatactcaaaggcaggaaataaggaaataatggcaaaattgtccgtaaatcaaacgggaatgtgtaaaagatgtgggtgatgaatgttatgtgct
atcaaaaacaattgatttatacgagtgaagtccccatgacttttcagctaagtgtctcagtctgtctactaaaatgcacaaatattcgcgatagaggagagatcgtaatatcaggaaattgggctggtcttatgTTG
ATATTCATTCCATTGTTTTTTTAAAGTAtgtttttgtttgaaactggaatttttcattatgctgagcaaacatgttgaataaattgacttaaaatgttgtgcgccaatgctaaaatgctaaattgaa
ttcacttaaaatcttgaattaaaagtctttacgtatatactgtataacacaaccatttttaaaagaagatatttttctaattttctcattgctcgtacttgcctaacatgtctaaataaaaaagaaacctttatcttattagtttgta

gccctgttctttgcttttatgcttaaggatgagaaattaaagctgagaccgcc
>TRINITY_DN33377_c2_g2_i1 EF1A
atctggtgatgctgctATGGTTGACATGATTCCTCAGAAACCAATGTGTGTAGAAGCCTTTTCAGAATACGCCCCGCTTGGACGTTTTGCT
GTCCGTGATATGAGGCAGACCGTGGCTGTTGGAGTCATTAAAAGCGTTGAGAAAGCAGAGGCCACTGGAGGCAAGGTCACCA
AAGCAGCCCAGAAAGCTCAGAATAAGAAGTGAttagacgttcaattagaaaacgtcaaatttgatacaaaaagcacttcaaattactgtagattttgaaacacattcacttactttata
aaatcaaagaccattttaccatcttgcattgtcccatcgcataaaatgttttacgaattattctggctatttagaatgaaaatgcacaaaaaagagagaaagaaaactgtggtctgagcccttttagaatcaaaTCT
GCTTTATGTTATTAAAATTAAAGTCaagtgc
>TRINITY_DN56443_c1_g2_i2 EMC7
caaatcgtttcgacttttacttttcacaaaaaATGGAGCTGAGAAGTGGACTACTCATTTTATTTACGTTTTTACTCGATATAAGTCGTATTATCGCTTC
AGATAATGTTATAGATGCAGATATTATTGATCGTTTCAAAATTGAAGGAAAAGTCGATCTCACAGTAGCGAAAGATAAAGACTGG
TTGTCTCAAACCAGAGTTATTATTGATGGTGGCGAGTTCTTGGGAAATCTTAGGAGTGATGGGTCGTTCGTGGTGAATGATCTCC
CCTCCGGTACCTATGTCGTGGAGGTGGCCCACCCAGAGTACATATACGAGGCAGCAAGGGTCGATATCACCTCTAAAGGAAAAA
TCAGGGCACGCAGAGTCAACTACCTACAGCCGTCTATTGTCAAAACCATACAGTATCCATTAGAGTTTAAGGAACGACAGAAAG
CAAAATATTTCCAGTTACGAGAGCAATGGAGAATTACAGATTTCCTAATGAATCCAATGGTACTGACAATGGTTTTGCCACTTTT
ACTGATCATGGTTTTACCAAAGTTAATGAATGCAGCTGACCCAGAGGCTCAGAAGGAAATGCAGTCACAGATGAAAGCTTTTA
ACGATAAATCCAATCTACCGGACATGTCAGAGATGCTGACTTCCTTATTTGGAGGTGATCAGAAGAAATCAAGTAAAACAAAAG
CTGTAAAGAAAAGGTGAtaacacggcatatgtattattgttaacattacacaggaaatatgatgaattatttattttcccttgtatgtttattcaaatacatgaacgtctctctgaaaagtcttaagacatt
aactattttatgactattttattcttgaaaataagttgaactttcaaaattgtgtattattgtcagtatcaaatttttgtttgctctattgttaattcaagcagggaataaaaccatgtgaaggaatgtttaatttgatagataatta
gaagataaatgctgctaacacaggaaagtgaaaaatggtcaagctattcatttcggtaataAATGCACAACTGAAAATATTTGTTAAAGTGgttatccaataaaagtagg
>TRINITY_DN46676_c0_g1_i3 ESI1L
gtcatcatttaactagttaaattcgttttggatgtcgcttcttttaatttcatttgagaggttgaaATGATGTATAATAACAACGGCTATAGACCAAGATACATGGGAGAAAT
GCCCAATCAAAATCACTCAGATCCTCCAGGAGACAGGGAGACTGAGCGTGAGAATCCCCGGGAATCTCAGCCGGCTACCCACA
GTACGGTGGGTTATACTGTTGTTACACCAAATCTACAGAAAAGGAACCAAGTAATGCGACAAGCTGAGCAAGGGGAGCAGTTT
TATCAAAGCTACAAAGATCAGCGTAAACTCGGATCAGTGAATTGTGGAGGTGTGCTTGGCGGCGGACATGTATCAATGCACGAC
GCCCGGCGGGATTCACAAAGTGCCCAGAGACGGGCACTCATAGAAAGAAGACAGAAAGAACAAGATCGTAAGGAGGCAGAG
AAAAGGAGGGAACAGGAGGAGATAGACAGAAAGAAAGCTGAACAGAGGAAGAAAGCAGAACAGAACAGAGAGAGGGAAC
AACAGAGAGAGGAGAGAAATCGTCTAGAACAGCTACAACGCAAGGAACAATGGTTGTCTAGGTTGGGAGGTGGAAATACTGG
TGTTCCTGCTACAATTGTCGATCAGAGACCTCATCTAGTGCCATGGAACTTGAAGAAAGTGAAGAATCACGGACAAGCAGCTAT
GCCATCAACAACATTACAATACAGAAGCGGCTTGGGGTGCTAGtcagagcctttccacattatggacaaacttatctccatgaacttctacagcaatgtaatggatccgca
gatgctgtgattgatattctggaataaacttgacttatgctaaaatacagagggtttccaacacaatgtatctaaaaccagaaatgaggacagttgtaatctagttatcctcaccattctctttcccattagccaatgac
ccaagttagttcttggagcttcttggattcattggtttacaaattgtctcttgcaaatattctgaattaattgaccttattcttaccttgactagagtatataaaagatgaagattttttttgcagaatacatgcagcatcatac
caaacgtccaaattcaatatgatggctttcaattgcactgtctgtcataaaaaggcatcaaatcacaatgttttaactgctacatgtatttaactaagtaacaaaagtcttaagaattttattcaaaacacttgtacgctca
ttaagcacatgttatatgtgtacactttaacatacatagtatgtttagaaagaaagaaagtgattctttacataaataaatttttacattaatagtagatatatgaaattttctgtgatttgtcttgcgttctgtgttatatgtaac
aaatactttaaacaagatagatctttaaacaagcaatgtgcgtatgaaacatctagtatATTTCAAAAGTTACAGCCAAGCTTAAAGTTttgaacaaacagactgacaagaaaataata
atttgtccaaaattttcaatacagaggcatagaataattcaagattactctgacaaacaaacatatccttggttttagacactatgcccaaatggtaataaactgctacataacataaaaacaaaaacaaaaaaaatat
aagcccctggtcattttaataaataaacttttcattggctagtctgaatgcaataaatgttgtcaatctctccgtg
>TRINITY_DN42991_c0_g1_i1 FHOD3
gcgtgtgttgaatcgttatcataaacttctgttgtatATGGGAATTCCTACAAACTTCGCCAAAGATCTTAAAGTGAATCATTTCTGTAAAACTGTGAGT
GAATTCGCTCTGGAGTATCGTACAACAAGAGAAAAGGTCCTTCAACAAATGCAGAAAAAGGCCATGCAAAGGGAACGAAAGA
AGACCAGGGGCAAAATCATTCTAGAGGCGGGTAATTTCTCAAGGAGTCGATCAAAGGACGATTCTACGGACCACGATGAATTG
TCAAAGCTCTTGTCCAATGGCTACACAAGCGCCGACGAGAGGGGGTTACCGGGCCAGAAAAACAGGAGTCGAAGGCCACTGG
ATTCTCGGCATTCCTCTTCCCGAGGCTGTATAACAACGGATTCCGAGATGTACGACACGGGAGAAGACGAGATTCTGGAGGCAT
GTGTTAGGTCAGCCACGGCAGCATCAACGAGGGCACCAAGGGAGAGAAGGAGGGCGCGGCACCATAGAAAATCATCAGTTTG
GGATAGTGATACTGATCATTACTAGtacgacgcaccctttaaagtggattgaactccgacgaga
>TRINITY_DN51861_c0_g1_i1 FRRS1
cactgattgacagacggtaagatgtcgtaaaatctttttgtcaaatatgatgacagtcgtctattgatggtccgttatgccctaaccacgtgacgtgaatccgtaattgggccccttccgtcgcttatttcccattgttttc
ctttgactttagactttttttctaatATGGCCGCTAGTTACCAGAAATTCACTGTAGCAGTGTTAGTTTTGGTTTCTGTGATTCTGTACGTAAATTCC
TTCATCATTGGAGCACCCCCGGACACTTGCGGCAATCCTACCACGTTTCACACGGAACCTTTAAATGATTCTCATGTTGGACAGT
ATCTACCACAAGACATAGACAAAGCACCATATAAACTCTTTGTTAATGCCAGGAGATATGAAAATTATACATTTGCGCGTCTGAA
AGGAAAGGAGAAAAACGCAGTTAGAATTACTCTCAGGGCTGCACCAGGAGATTATTTTAAGGGATTTTTCATTCAAGCCACAA
AATCAAACTATTCCCTGGACCGACCAGACAGGCCGGCTTATGGAACCTTCCGTCCGGTTGACCAGAACTCAGCACCTCGTTCTT
GTCGTGCGGCTGTCGGCGTCGTCGGCGGGATCACACATACGGGTAACACCAACAAAACGGAAGTGTCTGTTGATTGGATACCG
CCAGTCGGGTGTAACTTAGGGGATGTCCAGTTTGTTGCCACAGTTGTAAGGGGATACAGTGAGTACTGGGTGGATATTAGATCA
GATGTAATCCAGCCTTCAGGATTTGTACGAAGAAGTCTGCTTTGTGAATTGTACGTGGATCCATATAACAGACGATTACAGAGAC
GTGTTTTACAAGCATTGCGTGAATTGAGGAGACGTCAGAGGCAACGCGGGCCAAGAAGGGGTTAAttaaagtctgttgtcaacagaagacattg
gttacaagctaatcactatctgttaatcacgatctgtcgcacggggttcaaattaactttcatgaaatgaacagattttcaaaaaaaaatgcaaaagtcaattatttgttggtttctttcaccacagttttatgtagttttttt
gatatTTTGAAGTGTATTTGTATAATTTAAAGTAattcttctttttctaactccgtgctgacgattattacttctcctttttttatctctatttttcttcctttctgtgtcattaaattacatttaaattgtag
atctcgtttcgggaacttactgtttcctaaagtgctttgaaaacgcagaattatgtctgcacgatataaaaactaactttgactgattaaattcactagcgaattaaataacgaacatttattcgaataaagacaagaca

agacaagacaagacaagacaagacaagacaataaatagatagtaataaatttcacaataaagtgatttttgaagtcggttataagtgaacattgtattttcgaatatttgatatatacgcgaagtgtatttatgcgtac
ataagaaaaagtaagagaaaaaataccaaattgaaaataaaaattcgccattcatggtgatgtatgtacatgattcgacatttgattttgattacagctcgggaaatgtctacttgtaaatgaaataaaaaatgtattat
aacttaattagaaggataaactttctgaaatcgacgttaaacagtaaaatcttgttgaaatgagaattgcaacgcctaatgttgaccttattatattatctttacgcatgttatctttgcattcaataagatttgaactcgtga
aaattagagacagaatgtaattctttttaccctcaagtcaattgtattttacactcgatgaagtttttatgtcatacctaatttttgtaaacgatattttaggggatatctattgaaagttttgattaaaatgaaaaaggcaaac
ttgaagtgattttgttcatattatattcaaaagtatcttgtccatgtaagagctgaaattgttttgtattcttgtcataaaatgttataaattatttttatataaatcccgagtccaatctaattggttgaatgccatgtaccgctta
gagaattttcgccccttttaacatattggccattcgactggaaatcgaattaaaacgatataaaattgtatttttactttgcgaatgaaaattcaaagattcggcgaaactacttatcacaagaccaagccaaataaaat
actgtttacggtaatacatctcatataaaatacttgaagggtgatcaaaggatgagataagatgggaaatctcatatccttttcttataaattgtatctgagacttgacacatcacttggttgatgataagctcatgggc
ggatctagaaatttaaggggatc
>TRINITY_DN48744_c0_g1_i4 GOLI4
aaaaagaagaaattgtacattttattttcactgtcaataaggaaaaattaatcatattaccagctgacataaattttgtattgattttacctgttgtacaggtaaaaatttcccacaatgcaacagaaaacgacacgtcgt
cgccatattgttttgtttgggatgcaggaaacgtgtcaaatcgcgagctgaggttgtgttatcgccaatcttttccaATGTATGCACATTCTGACAGCAAAATGACTCGCAGAAA
CGGCTCTGTGATAAAGTACATTGTTTTTATTGTGTTATTTTTGGGTGTTGTGTACTTTATTTATATGCATAACAAAGTGAATGAAAA
ATTGAAGGACACAGAGAGAACTGCGGAGAGATACAGGCGAGAACAAGAGTCATGTTCTTCTCAATTACAAGATTTATCAAATG
ATAAAGAAGAATTAAGAACCAACTGTGATAGAGAAAGAAAAGAATTTGAAAATCGTTTAAGTTCGTTACAACAGCAACATAAA
ATGCTCAAAACGCAAAATGATGATTTGCAAGCAGAATATAGTAAATTAAATTTAGATCTTCAAAAGGACAAAGAGGAGCAAAA
ACAAGTAGAATCTCAGAGAAATCAAGAATATATGCAACTTAAACAAGAAAAAGAATTAGAACTTACTAGTTTAAAAGACCAATT
AGCCAATTTAGGTAGAGAAAAACAGCAATTAGCAGATGAAGTCAATAGTTTGAGAACGCAGTTACAAAATGCTCAGGTAAAAA
TGAATAACGAGCTACCCCACCCTAAACCCCCTCAGTAAccatggtatttcataacaataaaagttatgtacaacaacaaaaaataatcgtttacatgtactctagatgtagatgc
atggctcattcatttctagtcaactttttgtttgtagctagccattaaacaaagtttatgtagttgatgccgtgcagggcatatctaatactgattgcccatttgtaatcttgtgtcatgcttcttgtagaattctagcatcata
ctctatccttgcttcttaaaatgagaaatataggcccacaatatatgatttgaagatgaaaacttgcctaatagaattttttttgtcttacgtttcccaaatgtggaatgggtatacagaactaCCAAGGGCATA
CTAGTTATGTTAAAGTGcttattgatgtatgtttgaaacatatagtgatgctactgccttctttataagtggcgaaagaacaagtagaatttatgtgtcctggaatttcagttatgtacatgtaattgtataa
attgggggaaaaggggggagaaagtgaaaagagaaagagatggaggatgggcctcatttactaaagaataaggtcattgaagggattgccgagtgtctgatatcacacattctgattggcttttatattgaga
catcataatcaagtatcgagtaaaaggattctttgataggccattcaataatgacgtcaccatatatacatactgtaattctgccaacatttgtaatgtctgatttccgctatctctcaatttctagtagatgcctattatgct
aattaatattatctatgttgttttcctttatttattttgtagaatatgttattataatgcttttgttgtaaactgtctcttctttttatgcatgtgaattttgtgcatttatgtttgaaaaaaaatctcgttttgctttgtctaataaatctga
tctagtcaca
>TRINITY_DN50951_c0_g1_i14 KAT3
actgtgtcatcggtgagtgcggggctagtacacgaatttaacactttattgggtttcatcaaagagacgaaATGTCCAACGTATCACAGAAACATGGCATTGCTGACAGAA
TCAAAGTGAAAAGCAAAACAAACGTATGGCCAGAGATACTGGGGCTGACGGCAAAACATGGTGCCTTAAACCTTGCTCTGGGA
TTTCCTGACTTTGAACCACCGACGTACATCAGAGAAGCCTTGGCAAGTGCGGCTACTGGCAAAAACTACATGTTGAACCACTAT
ACACGCGGAGCGGGTCACCCTCGTCTTGTGAACGTTCTGGCCAAGCTATATAGCAAACAGTTACAACATGAAATCCACCCAATG
ACAAACATCCTTACATCTATAGGAGCTTTTGATGCCTTATATACTACATTCCAAGCTCTCATAAACCCCGGTGATGAGGTTATACTT
TTGGAGCCATATTACGAACCTTACAAAGAGATTTGCCGTCTTGCCGGCGCAAAAGTAGTCTATGTGCCATTAGCACCGAAAGAG
GACGACTCAAACAGCGATGGCTGGGTTTATGACCTTGAGCAACTACAGGCAGCATTCAGTCATAACACAAAAGCGATAATCGTC
AATAATCCCATGAATCCATTGGGAAAAGTAAGTTACTACAAGGAAGCTTTGACAATTTTTTGAtgaactaagcggaaatatttcgcgacatgcatcgg
atataaggagcacgtatacttttatggtatagataaaccaaaattgtcttcaaaatgaagcttttgtcatgataatataattaatgtctttgtttattatattcctaatttcccatttctcactatttctttctttttaactacacaag
gtctttacaaaggatgagttgcagccaatagcagagctttgtgtgaagtacaatgttctatgtgtggccgatgaagtgtatgaataccttatctacggtggtaggaagcagtacagaatgggTAATTGTGT
AAATTTATCTCTTGTTAAAGTActgagcatatcaacgtgatcctacgatatttacggaagaacagtcacaatgtgaaaattatgtaagttatagatattcatggtcgacgttcatgttacttgtatgt
tctcaatataatgtaactaaattattacatatagtttgttgtacatgcagatatgaaaaaataaaattattttcaaattttcataactatttattaaagaaaactttttttccagctaccttccctggtatgtttgagcgcactataa
caataggtagtgcaggtaaaacgttcacggtgacgggatggaaggtgggctgggctataggtccagaaaacctaatagctccactacaagctttccatcataacatcatacgaaccaaccccacaccgttaca
ggaggctgtagcggtgggatttgagacagagtttgagaggttagataaacctgactgttatttcacgacattgtcacaggagatggaagagaagagggatcaactagcgggatatctacgtgaggcgggacta
aagcctatcataccaggaggaggatactttatcattgctgatatatcatcactagagatggatcacaattacttaaatcacgatgaaagctacgactatggtttcgtgaaatgggcgattcaacacaagaaattttgt
gcagttccaatgtcgattttcttcagtgatgaacacagacatgtaggagaaaaatacgtcagaatctgttttgttaaaaaagaggagacaatgcaaaaagccaggactatcctacaagaatggaataccataggc
aacggttacagtaatgggcatagcaatggttacattaagggacattccaatggttacagtaatggtcacaccaatggttatataaatggacactccaatggacatgtagatgcccctactgggattagataattgag
aaaatatgtgacaagatgcatatagaataactatgaaattaacatttacgcttattgaaggatatatgtatacttcaaatcgattttgtgtgcaataatcctgttcttttaaagattttcaaattatcaggcttgaaagaatag
aatcttttccagtatcatatatatatacatgtatgtatgtgtttgttaattttcattaatttttttcatgattttgcatttccaatatacaatttttgcattaaatttattattatcaattttatctaattaaaaatgtaactaaaatcatgtaa
atgttgttaatgaaatcaatacaaaaatgtatttctaaatcatcagctaccaagagtattaaagagtcaatagaatgcattaaaataatttagatattacatgttctagaatattgctggctgcccctaatgtttgagccta
attgtatggcaatgttgactcttgttgtgaataaaaatgcaagacattgacacagttgttagcatttgaaaaaaaaaaaaaacttgacctaaaagatgaccttgtttgtgtatcgacattctgcaatttgcaagtcctgga
actttcta
>TRINITY_DN52847_c0_g1_i2 NAC2
tgggatttttgtttattttactatgattactttttttttcgactttggtttttgtggggtttaccaaagatctatcgttccagaaatgtttctagtttagcatccatgatatattattcaaggcatgattcgaaagtttgtgtgaaatt
cactgatagaaaggatataaaatatgtaagggactgtaactgaaaatttattcaataatttaataacatgattaacatatatttaaagtgacattccatctaaagagtaaggaatcagcctcaagtaactctctctcgtta
atttgttgtactaaatgttctttttagctattcaggtagtggacctcttcttcacatatatattttgattttctatatcccactattgcaaacaggggaaaataattggtttgctcttttctgaattttcaaatacatgtatataaaa
agacggcaccatccaccactaaattgatatactaactattgattatgtactgatctgtcatctttaacatttcctctcagctctatgacacagtgactttacccgaggaggaaagaaccgagaataaactgacggtga
aaaaggatcataccgagggcggcaccaaggccgtggtagaattcactgccgcctcactctcggttctagaaaacgagcaaaaggtccgcgttggaatccgacgatatggaaaactggataaacctgtgaaa
gtcaaagtagagaccataaacggtacagccgtagcccctgaggactacatccatctagaggaggtgattgaattcgccgccaatgaacagctacgacagttgtacatacagattgtggatgataatgaatggg
aaccagacgagttcttctttgtcaagctagccctgacacacgacgaggaaactaacgagagcgtagtgctaggaccagtgtcaatttcacaaatcacaatcattaatgatgacgaacccggaaaacttgaattcg
ctaagccaagtatcatcgccaaggaaagcggacgtcatgtgagaattcctgtaaaccgtatccatggtgccgatggtcacgtgtctgtcaagtggagaaccaaggacatcactgctatcagtggtgtggattac
gaaggaggagagggagagctgaagtttgaccatggcgaaactaccaaaaatatcgacattcatatccatgacactaataaagatgagagagatgaaagttttgaagtagagttaggtgaaactgacggagga

gcagagctagccaagatcaagaaaactattgtcaccatcgttaatgatgaagaatttaacggactggtcagcaggattgtcaatcttaccaaagccaacttagacgccctacaactggaggactcgacatgggt
cagtcagtttaagacagccATGAACGTTAATGGCGGAGATCTGGAGACGGCCACTTTTGTGGACTACATTTTACATTTTTGCACGTTCTTTT
GGAAGATTCTGTTTGCCTTTGTGCCACCACCCAAATACCTAGGCGGTTGGCCAGCCTTCATTCTATCCCTCGCTGTGATTGGTCT
AATGACGGCGCTCATTGGAGATGCAGCTAGCATATTCGGTTGTTTGATTGGTCTAAAAGACTCCATCACCGCCATTTCTATCGTT
GCCTTGGGAACCAGTATGCCGGATACGTTCGCCAGTAAAACAGCGGCCGTGAACGAGAAGACAGCGGATAGCTCTATTGGTAA
CATCAACGGAAGTAACTCCGTCAATGTATTCCTCGGGCTGGGTCTGTCGTGGCTTATAGCGACTATTTATCATCTAGTAAAAACT
GGTTCGGATTTTGAAGTGCCAGCAAAGACTCTTGCTTTCTCTGTTGTTCTCTACACGGTGGCAGCCATTGTAGCAATAGTGCTTT
TGGTTCTTAGAAGGACATTACCCATATTTGGTAATGCCGAACTCGGCGGACCTAAAGTTCCTAAAATATTCTGTAGTATCATATTC
TTCTCACTGTGGCTGTTTTATTTATTGATGTCCATTTTAAAAGCTTACGATATCATTACTGTGAATATTTAAttgattttattatgtagatactcaatga
tacgtaatagtaccagacatacggtatggaagtacagtacaataacaatggttgttttcaaagatatgcatgcgcggatctagaaaagtatttttctgggagggagggaagcagagagggagaggggagggca
aacgtatgacttttcaaaaaaaaaaaacatatacgctaaaggagaaaaatggaaagtaagcatttatgacaatttccaggggtccatcctttcgatccgcgcttgatatatttgaatttattttgttgtaacatcgttatg
aatttgaatgaatgtttattcattttgaaattcttatatccatgcaataaggcattgaaaacttagggtagctacattattgcttgtctcttataacttttacatttcctcaaaaggaaacattttgacagtttaaacgtatgtcg
tttacgtacaagatctagttgatttcgttttcaaatgggtaatttctgaattcatttgttaattttgagtcgggtaaaagaaattagaagagtgactaatggatataaggttaatttgatttgacatataaacaaagattttaa
aggtttttgaaaaacgtggctttgaaaaatccattacttatgcattattttcatcgctagaagatatgcacaatttcagaagaattcgagcttgccatttgtgccattctaaattatataccttgccagttcatcatcataatt
actatcacacaaattgtacctgatacatcatacaagcttttctgttagaacacttcattagtgcgattccattccaattgtttcgtgtaaatacattttagaagtcttcttgaaccggatttaatttttgaaagacttaaaattt
gaatttagaataatagatctacaggaaaaatccctttgaacgcttttgctttccaaatctgtaaccaaagcacaatactaatttcacagatttttacacaattttttccaacgataagttatgacttcgcgagtctgctttca
atctttggtaattaatattagaaagtcagttaagaacccgtactgtttgtagattttctgtgcaaatatttgtagtgagcaaagcttggtctgagaagaacattgtatacatgtagtttaattatccaaatatttatcacacc
ggattacatacgatatttgaattaaatccaatggaaatgaccttgcttaaattcatcttcagattatgttctctgtattaattatgtcttttgggtatttcttgtaccttgttttgtactacgcgttacattatatataatatctcttat
tTCATCATGCCTATTTTAAATACCATTTAAAGTTgttgtacattttaatattgtattttattcttttacatatctatgttcttgttatatttttagtttaatttctacttattttactctatttttatattactta
tacattattgggctgtagacactacatgtatttatatgtgccttctagaacgacatgccatgtatgtgcgcatgcgtgttataggctggcatctctaaagggtctctctttgtttgtatattttgctttttaataataaatcctg
taaattttatgtacaaaaataaaactttaaagccgtaaaaaaaaaaaa
>TRINITY_DN58441_c2_g1_i4 NBR1
agcctttagtcaacaggatcacaccagtcaaagaagttcatcttgcagaggtcaaggacacaagtgttgactcccaggtcaaggacacaagtgttgactccaaagtaaaggcagtagatgaaaaagtggaagt
gtttttactggagaaaccactcaacaaagactgtagcaacgaattacgggaaatttccatcagaaaaacggacaatattgaaaaatctgaagctccaaaatcagaagaagacatttcgtcaaagatagcagagg
aaactccatccgaggaagaggaaattccagaagtgatcgatgattgcgtacagATGGACAATGATTCATCACGGCTCGAGCAGGATCTTAATTGTGCCGTTGCC
GCGGTGGAGAAATTGAAATTGGAGGAAGAGCAACAAGAAGCTACAGTTGCAGGACAGGACCTGATTTCTTTTGATATGCTCGG
AGGACTCGTTGCTAGGGGACCCTCTATGACATCACACACGGCGACTCCTAACAACACACCACTAGTGGTCTCCCCTCCCAAATC
TCCGGCACCGGAGCTGGATAACGATAAAGATGAAACGTTGTCCACAAGCAGTGTAGAAATCATGAACGCTGACATGCCGGAAG
AAGAAGAAGCTGTCCTGATCGACGATCCTTTTACCAAAGATAATCTGTGTAATATACAAGTGTCACCTCTAAATCGTCAAGCTGA
TGAGCTGAGATCTTGGAGCCCTGACATTGATGAGTTTAGCACAGACGATGAGAGCCTAAGCGAGGACGATGACTTTGTCTTAGT
ACCAATGCCTGATTGCTTTGACACAAGTAAACCAATCAAATCGTCTGTTATATTGGATGATTGGTCAGGCTCTGAGGATGAGGCA
CCCCATGGGGACGTGGAAACGGACGAGGCTCCAAGGGGGAAAAAGGAAGATTTACTCAGCACTCAAAGTAGCAAAAGTAGTG
AAGATGAATTTCGTCAAGTTTCTGTGGATGAGATCTTGACCGCGTCTGGATCCCTCTCCACCCCGCCACTGGATCCTGTAGTGAA
TCCGCCCCTGGCTGAGCCCCAGATCACCATCAGTGAACCCGCCCCTGCCACTGTTGCTATGGTAACACCTATACCAGACATACA
GACTGTACCAGCGGAGGGAGAATTCTTTGAAGCTTTGTCTGAGGTACCAAGAGGGGAACAGGATATCGGAGCCTCAGGAAATG
TTGACAGTCCACCCCTCAACACAGAAACAGGGGAGAGAAGTGCTAATAACACAGAAATCCTCGCACAGACGTCACCTGATAA
CGCAGGGGAGAGTACTCACGTCAAACATCCTCTACCCTCAGATAACGCCCCTGAAAATCCTAGCGAATCCACCACTGATAATAC
ACAGGATCTCCCACAAGGTGCCACTGCAGGTGACAATCACAATGCCAGTGAGCTTGTCAATAATGTGATAAAGGCTGCCAACC
AAGTGGCATCTAACGCTTACACGACATGTAAAGAAGTGTTCTACACATGGCAGGCAAGGACTTATCAGAATGGAACTCCAAAC
AAGTACAAACCCCCACAATCAAATTGGAAGCCCAAGGAGGGTGCATGTGCCCCACCCCCACAGACTAAATGGAAACCAAAGG
AGGATACGTTTGCACCCCCAAAATCTACATGGAAACCCCCGGCAGAAACTTGGACCCCACCTGTCGAGGAATCCCCAAAGAAA
CCAGAAACTGGGCCCATGGGCAAGTTGATAGAAATGGGATTCTGTGATAGGGCTAAAAATAAGGTTCTCCTTGAAAAACACAA
ATATGATGTGGAGGAGGTGCTTGGAGAACTGTTAAGAGACATGGACAACTATTGGATGGATAATAGGCACTGAaaatgttacaaagaatct
gattggttgaattttgtttgtgatgtacagatgtgaattacttttggttaattttcaatccttaatcctttgaaatatatttaaaactcactgattggctgatttggatcacagcaaaaaaagaatatgttaaaaaatcactgat
aggctgctttcctttagtgtgtaaggatcacagataaaaagaaatgttaaaaactcacggattggctgatttcctttagtgtgtaaggattgcagaatttttttgattggctgaacaagtttaaagatatatgtatatatact
agatatgaagaaaaaacgttatattcttcacgaacactatctttgattagtggatacttaaagaaaatctcttgacatttttttgttatatggtatgaaaaatttacattataaccatgcaatttatcctcatacaatttatcctc
ccccccacacacacctatcgattagctttttccactcttttgaaaaaataaaagcactgcagttttgccatcacttctggttaagggctcattcacaaatccattttatgtatcttatacaatatgtgtacaatttgataacc
agggctttatttatttcataatgcaagttttgaatcaactgcagTTGTCCATTTATGCTTATCTCACTTTAAAGTAcagaaaaatttctattgtctgtgaaatacaaggtatgttatgtagtg
ggatggaaaatgtccgattttcaatatgggattgaggctcgtcaaattatggctgttagaaaaaagttaaatatggtggacattttcgcgcttctttggagataggattagaaattaaagttaacgtctagaaacttcac
atcataaagctaagtttattgaattaaagataagattatattaggatatatatataaggggaaagag
>TRINITY_DN52868_c0_g1_i4 PA2HB
tgaacatgacgtcacgaccgggaagaatttacaacttaacaacgccgtgatccacttaggaaaacctgaataggtgggacgttcatatgagaagtttaacatcatacaggtgacaaaaagcATGATACA
GAATGGACTAGTGAGGTGTGCACTCATCTTTTACTGTTGGCTCCAGCTAACACAGTGTGCATCCCTAGAGAGAGAAAAGAGGG
ACTTATTCCAACTTGAGGAGGTGATTTATGATGGAACGGGACTGAACGCTCTCACCTACCTTAATGGTTACGGTTGTTACTGTGG
GTGGGGTGGGAGTGGTGCCCCTGTCGATGGTATAGATGCCTGCTGTATGGCGCATGATAATTGCTATGGCAACGTAACAGCCAA
ATCTAACAACCAATGTACACCAAAATGGATGCACTACGTATTCTCCATGAAGAAGAATCAAGCAGGTCCCGTCCATAATATTGTA
GAGTGCCTTGACAAGAAAACCCATAAATTTCCGAGTACACCGGACTTCCAAAACCCATACTACTGTCATTATCAGATCTGCTTGT
GTGACAAAGGGCTGGCCGAATGTGTGGAAAAACATAGTAACATTTATAAAACGCACCCGAAAATGGATTGTAAGAATCCGTAAa

gtgctcgccaagtttgtctatttacaagagagagttacctccccatttgtgcatacacgtttatgcaatatagacctctataatactgtcaaactagatgtggggaaacatttcgtgcagagattgcagacaatttttgtt
ataccagaaagggggggggggggaagcaaaggctgtctgaaatttggaaaaaatttaaatctaaaaagggagagcaatatgaatgtgtagcaaaatgaaattctttacaaaagacccgatatatcttgagtcttaa
gtttggtttagtttcataaagtgcttgcgtaaaatcttatcttctgcgtaaatcaatctatctattgacctaattagcattgtcttccatgaatatttattgacatatcagatatcaactttatgccaaatttagcttaaatcagct
ttcttggatacattcgacaaaggttttcatttgtattttcctcattcaaaaaagatgaaaagggctgtgccatttcattgttttgaatggagaaaaattgtaaaagagatttgagggtgaagattattttttcaaaattaagt
ctgtatcggatattgacctcaaaagggaagtaACCTCAAGGAATCTCTGATTTTAAAGTGcttaatgattctacatatatttattttataaattgattttagtcatgttagttgctgtgtactttat
caacaataacgaaaatctttaaggatcacaagttcaatcacaagtttaaatcatattaaaagtgctaaataattctacacatatttattttattgcttttagatgctgtgtatctaatcattgtcagattatgaaagactaaag
aatcacacgtttaaatcattttcaagtgctaaataattctacacgtatttattttat
>TRINITY_DN55201_c1_g1_i2 SRS10
agtgcgtcagcgcggttcgaaacattagacggtcgaaataacttcaaagttcgtttgcttgtcgcagatcgcaacgacgcgATGTCACGGTATTCCAGGCCCCCAAATTCATCAC
TTTACGTTAGAAATGTTCCGGATGATCTTAGAGCCGAGTCAATGCAAGAAGAGCTCAGGTCATTGTTTGGAAAATATGGACGAA
TGACAGATGTTTATGTACCTGTGGATTATTACTCACGAAATCCCAGAGACAGAGCAAAGCAAAGAAAGTCTTAAgagtctaaggtcaaaa
gtgtacaaagacaagcttctcaagaaagtcaaagaaatggttggaAGAACATTTACAGGCTGAAGGTTAAAGTCacagtgttaaaagtcaaagctcacaaggaaaagttaaagac
ac
>TRINITY_DN35472_c1_g1_i2 SYWC
atacttcagaATGACTCGAGATGTTGCACCAAGGATTGGTTATCCAAAACCAGCATTGATTCACTCTACATTCTTTCCCGCTTTGCAAG
GTGCTCAGAGCAAGATGAGTGCAAGCGACCCAAATTCCTCCATCTTCCTTACAGATTCTGACAATGATATCAAAACAAAGATAA
ACAAGTATGCGTTTTCTGGGGGCAGAGCAACAGAGGAAGAACATCGTGAAAAGGGAGGTAACTGTGAGGTAGACGTGTCATAT
CAGTACTTGACATTCTTTCTAGAAGATGACGAAAAACTGGAAGAACTAAGAGTCGGATATGAGAACGGCAAAATCCTAACGGG
TGAGCTGAAAAGAGAACTCATAAAAGTTCTTCAAAAAATAGTTGGAGAACATAGAGAGAGACGTTTAAAAGTAACGGAAGAG
ATGGTGAATGCATACATGAAGCCTAGAAAGCTTGACTTTGAATTTTGAtttaacgccatcttttgcatactttcatcgtattgctgtcatacatcaaaagggtacgtcatt
gtaacaacattgtgctaatacaaaacaatattcatttgtaccgtaaccactcctaaaacccactggaaggagcaaactatcatataaacattctgaagattcaataaagttttggaatgtctctaacatagactAAA
GGAATATTTTATTTCCTTTTAAAGTTctaaatgtcggttac
>TRINITY_DN53558_c0_g1_i9 TM87A
agacgtcccagttgatttacatgccttttacttgctaaggtattccccatgcgcaaatgcatttaacggcattgagcgagcaaaagagtacctgataagtccaacgatacatcaattaggagactatacatcggatat
catttcaaattcgacaaatgtgaaatgtgagaagggaaaacgtataacactgtccaattcgcaaaacgaatatttctccgcgtggaaggacggagcatatggattgttccttcaaATGACAAAAATTC
CGAATAATCTGAGTATCAAAGCGGAAATTCAGCTATACAGTGACCATGGTTATCTATCAGCAGAAGAATACCCCATGCTTATTTTG
TACATGGTGTTGGTAATATTCAACGTTGCGTTGTGTTTGGTTTGGAGTGTGGTTATGATCAGGGACAGGAGGGAATTACTACAAA
TTCAACATTGGATCACTGCCTATATTGTTCTGCGTACTTTACAACAAATATCAGCATTTAGCGTATTGAAGATCGAACAGACTACA
GGCTTTAAATCATCCCTTCCCGCCTTGATTGACACCTGTCTAATGGGTTTGAATAACGGTGTAAGCATAGTCTTAATCATGATGAT
AAGCTACGGTTTTTATTCATTTCGCCATAGACTTGGAAAGACGTACTGTGTGATTTTACTCGTGGCGTTTCTGAGCTCACTTTCGT
TAACCATGCTGGTCATTCCACAAAGATTACAGGTTAGGATGTGTCTACTACAGCAGTGAattatcatgacatgttttaaagttaaaatatttaaggtgggtcc
cgactcgaaccgtcataggttttcgtaatacgca
>TRINITY_DN47876_c0_g1_i2 TYR1
ctttctcacgtcaaattttcccccttacctgtacggtggttaacgctgccttagtgacggcatgggttcttaaggaatattacctatttttgtctctctagaatttttctacttttctgtaggattttgggacagttagaggaa
atttttctaaaattggaatagttcttaagagaagacaatactgaaaacATGTCGAAAATGCCGTCACTCCTCCATATTTTATTACTGATCATCGCCATCTGTCCT
CTTTGTCGTCTGCAACAAAGATCAGTGAACAATCTAAATCACAGATTTGTAACGTGGATGAATTCCTTATTTTATATACCAAACG
ACAGGAATTTACGAGTTCGTAAAGAATACCGGATGTTAAGTGATACGGAGAGGAGGGATTATAATAGAGCTATCCTTTTGCTGAA
AAATGATAGAACTGTATCTCCAAATAAATATGATGCATTAGCGTCATTACATCATCTTAATTCTGCAAATGGTGCTCATGGTGGAC
CAGGGTTCCTTGGATGGCATCGTGTTTATCTTGTGTTGTTTGAAAATGCATTGAGAGAAAAGGTTCCAAATATCACGATACCTTA
CTGGGATAGCACAATGGACAGTGATCTTCCAGATCCAAGACGATCCATCATTTGGTCTCCTCTCTTTCTTGGAAACGGCAACGG
ACCTGTAATGAATGGACCCTTCAGGAGGTGGAGCACGCCCTATGGTCCCCTCAGAAGGGACATCGGTGCAGACCGACGACTCA
TGACTCGACAGGACGTTCAAAATGTATTTTCAAGAAGATGGTTGTGGGAAATAACAAATCCTAGTGCTCGCGATGAGTACAATA
TAGAGCTTCTTCACAACCATGTTCATGTCTGGGTTGGTGAGCAGATGAGTAGAATAGAGTCGTCATCTTATGATCCAGCTTTCTT
CGCACATCACGCATTCATTGATTGTCTATGGGAGGAATTTAGGCAACGTCAGCGGCAACAGGGAATCAATCCAGCAAGGGATTA
CCCACGTATTGTAGGTGACCAGAACCATCAACCTCTGGTTTCGATGGGATTGGGAAGACTTCTAGTTATCGATGGAATAAACGA
CTTTTTCACCAGACAAATTTTCAGATATGAACGTCGTCCAACTTGTGTAAGGGGATCCAATACATGTGGGTCCCCTTACCTGAGA
TGTAACTGGTCTACCCAGACATGTTTGCCTTTAATAATGAGTAACAGAGGGACGCAGACGACAAGAAGAGTTGTTCAGAACAG
ACGACAACCTTGGTGGAGGAGATTCGTCAACCAAAGGCAAACATTCTTCGGTTGAtgaattatggataaagattattggattcaattccttgtgccaattcatg
gtttggggcttcttcattgtctttcggctttttgagaatttttgcatacgaacatcatttaaaaaaatgttgtctttcagacaaaagaaaaatataaatattacagtcatcattcattctgaaaaaaaatgtgataataaaaag
aaatgcagaatagaagcaaaaatgtttacggtacagtcggataatattgaacaattttaagaagacgttcctttgagcatgttgaagaaaagatttgaagacacaagttgtatgggattgcatggtagaatgtcttta
atcttgtaagttcgaacgcagaaatttcatgaggaaatgcatatgtagtgattgttattatgtataataatgcaatcctgataaaattgattttaatttatgttctatgtttcaattttctttgtcacttataggaggcattttgaa
aggattttattcaattcgaacatgttgtcaatgagttgttcattttgcatcttgatgcactcgccataagtaggtttccaaaatgccatttatatgtcgtttTGCTCTAAGTCTTTTTGTTTTTTAAA
GTAaaaacatattacgtattttctttataaaaggagttcagaagaatgctttcgattccaaaatatcttaaaatcatttaatatatgatacgtgaagtttggcaatggcacagctttatgacctcaaatattacaatctat
gcaatactaccaaagattcgctcgctatttctgcttcttttagaatgttttgttatatctcaccctaagtgtcttatttattatgttaagtaacaacttttaaaataaaatataaaactctgaaactttaaatgggagagatcaat
gacaaaatcttttgaacttaaaaattgcagttttcacctatttcaatgtaagaacttaatgaccatcccttatatcttttagatataatatgtacatacaaaataggatcttgtactgatctcaacaattgattgtaccaatgtg
aaccttaaatctagtctgtgtactcttaaggttgatttatactgcacaacgttattgtaccgtttaacaagttcagactattgatttttaagttcagtttcttgtcactcagcgatgtaaatacattagtaaaatggaaatccta
tataagacgtcaaagtgactaggtgaacatgattttattcgttttgtttttcacaaattgcacattcctatttattgagaataaactgttgttacaaaatgaaaaaaaaaagatcggaaga
>TRINITY_DN35658_c0_g2_i1 TYR2

cttacattatttttcagttttatcgattagcaagATGAAGAAATTGTTGGCGCTTGCTGCGAGCCTTCCCTTATTGTTATGCGTCCATTGTATCAAAGAA
AAAGAAATACTGAAGGAATCATATAAGCAAAAATGTATGAAGAATGCGGTCTATGACTTCAATTCTACAAATCCTACTACCCTGG
AGCCAAAATGTGCTACACTCTTTGGGCACGAGTACTCGGATATAAAGAACTTCCTCAAGTTCGACGATCAACAAATGAACTATA
TTTTGTCATTGGAGAGAGCTATGATGCGTACGCAACACAGGAACAATAAACGGCATAAGCGACAAGCAATGATGAGGCCCCGG
CAGGAATGTAGGACCTTGAGTGACCCTGATAGAAACGCTCTTTTTGGCGCCATCGTTACTTTAAAACAACCGTTTAGTGGCATG
TCCCGATACAATACATTGGCTGCAATGCACAATTTACAAGCATTTGGAAATGCCCACAATGGCCCAAATTTCCTTGGGTGGCACA
GAGTTTACCTGAATATGTATGAAGAAGCGTTACAGGAAATTCGTCCCGGTGTCGCATTGTGTTATTGGGATTCCACGTTAGATTAT
TTGATGCCAGGGGATAGTCAAAGAAGGACCGTGGCTTTCTCTGATGAACTATTTGGAAATGGACGTGGTGCAGTTATAAATAGT
CAATTTGCAAATTGGCGTCTTTCTGATAATACCCCCTTAAGACGAATGATCGGAGAGAATAATTCAAGTTTGACACGACCTGGCA
TTGTTGACCTTATTCTAACAGATCCAAGAATAAATCGTCATCGATGGATTGTGAATGAGGGATCGAGGTTTAATCAAAGTCCCAG
ATTTGGTTTCATCGACCCTGATAGCGGCATGAGACATAGCTGGGAAAGAGAACATGATAATACACATGTGTGGGTAGGCGGAAT
CATGGTTAATGTGGAGAGGTCACCTGAAGATCCAGTGTTTTGGTTCCATCATTTATATATCGACTACGTTTGGGAACTTTTCCGAA
GAAAAATTGATCCAATGGACAGATTCGATTTGCGTACAGATTACCCCATGGATTCAGTTAATGAGCAGCATAGAGCTTTTCAAAC
AATGGCAGGCTTTCCTGCTTACAGAAATATCGACGGATACCACAATTTCTTCAGAAGAATGTATGCACCTCATCCTAGATGCAGT
AACAACTGTGGTGGCTCAAGATTTCTAAGGTGTCCTGACATTGGTCCAATGGGAAATCCAGATCGTCGATGTGTATCTCTTGCAA
TCGATTCTGATGTGGTGCCAGCAGCGGCGGCTTCCCCTGCTGCAGCAATGGCTGGCTTTGGAGCATCTCGTGCAGGATTTGCAG
CCTTTGGAGGACCAGCAGCGATGGCTTCCGGTGGTGCCGCTAGAGTTTCTCTGCAAGCCACAGATGAAGTTGCGATGCGAGCA
GCAATGAGTGCACCACAAACAGTTGTCCAAGGTCCAAACTTCACTTCCTCGAGAGCTGATAGCAGACTTTTATAAagctaaaaattctttt
gctattcaaggttttaatggtgatttggaagtgaatatgtatgattctgtaattacgctggattctcaaaaaagtaaagatttaacgttcgttaatagataatgtgaaacgtatttaaaaatctcaagttcacccaatatatt
atttgaatgttttacagccttgctttcttgattgagggtaatgatacaataagagtgcgagtatctataattcctTTAGGCTCGAAACAGTAGCATAGTTAAAGTTtatgaccaaaatagc
acgggagggattgtgctagaattttaacctcctttttgtggacggaatgacaagtttccattccataacgcatccatacttgaaattgtcaattcttatacggaaaacattataatactgacattgaaatcgtagctttcat
tcttatgtcaattgtttctatcacacattcattgttgaaatatgaataaacatgattatgaagaccac
>TRINITY_DN57239_c0_g2_i2 VINC
catgtatgtaaaatatcattgttgtatttgtagatggtggtgccccaccccgtccacctctcccagaggactctactccacccagaccaccacccccagagacagatgatgaagatgaggctgccttccccaccc
ctcaggcaaatcagccaatcATGATGGCAGCACATGCCTTACATATGGAGGCCAAGCAGTGGTCGAGTAAAGACAATGATATTATTGCCGCA
GCTAAAAAGATGGCGCTTCTCATGGCAAAACTCAGTCAACTTGTCAGAGGAGAAGGTGGTACCAAGAAAGATCTGATTTCTAC
AGCCAAAATGATCGCTGAAGCTTCTGAAGAAGTCACTCGACTTGCCAAAAAACTCGCTGCAGAATGTACAGATAAAAAGATGA
GAACAAACCTTCTGCAAGTATGTGAGAGGATTCCTACCATTGGTACACAGCTAAAGATCCTGTCTACAGTCAAAGCCACCATGT
TAGGAGCCCAAGAGCCCATACCAGCCCCTGATGGCAGTGAAATTGCTTGTGGTTCAGAAGAGGACCAAGAAGCGACAGAGAT
GTTAGTAGGTAACGCCCAGAATTTGATGCAGGCCGTCAAGGAGACTGTGAGAGCTGCCGAAGCTGCGTCCATCAAGATCCGCG
TGGATTCCGGTTATACAATCCGCTGGCTGAGGCGCCGTCCCTGGTACACATCATGAgatgtgccgttaagatcgttatgacaggcttatgttagatagggaca
acaaaaaagttttataaaattgtacgtgcaatgatttatatatggtatcgaatgatgatctgaaatgagctccctttttgaggaagttcattatgatatagtagtgtctaaaatatatattatgttgctttttgaaattgcttcta
tatagcccgtagacaattatgaatgatttttttgtctcataatttcaaatgaatttatgtttaggtcatgtgtcaatcataccggtatagcatttatacatatacatgtaaaactaatgataatttattagccataattagtacat
gttttattaattgatctatgttcaaaattaaaaaaaaaaaatcttttgctttttacaaatatttaatgctgcattcagccaataattttacacctagaaaaaaaaagactatacatgtatgtacgtataggacttcgtgcttatttt
gacgttgatacattgtttctgtttttcttaacataattatgtattgtttgaaaatcttgttaaataagataacctgattagtagttaaaattttcatgaaacacatacatgtatgacaaataaaaaccattgtaaatttggccaa
atttgaaatacttgtaaattaaagattaattatttcgatttgacaacttccttatcatatcatcatttacatgaaaacatttgaagttacagtatttatgatagtggtgctattcttggtttgtaatgcacattttgatgttgatatt
gttcacttaatattatatattttgtagagtgtattttttttttgcattattacgtgtatgtaagataccattctttgttgataagaaatgggtatttgtaatcgacttttacgtaggcttttatatgaactacctttaagctatcattga
aaggcatatctgtttttataacgataattaatttaacatcgttaagcaatgatttctagatatttattgaggcttcgtgtaatgaatgcataaggaaagtgtgtagggaaattaatttgcaaaatgttcaaacatacacaag
ttacgtgtgtcaaaaaattgttatgtcatgacagatttttaatcaatattttttcagatatctataacatttagtgtgatataatattgtatctttattattatatatatatatatattgtgcatgctgaaaccctgtgtgacaccat
agtagactgtgatatacctctgtattttacaacccaagtcattctacctttgggaatatatcttaactatgaaaagccatataaattgatatgcaaaatttttatgtttgatgtcgtaattgatgcattcatactgacaaatcg
ttacacagtatggtgaacaaaggaagtcagttattAGATAAAAATAAGTGTGTTATTTAAAGTTaccaaggtatatcttgtttcagttttgtgtgtattacatggcaattacttataggaaaa
aaaggcaagacaaaggccctcttaatgctagcttatccttaatttgtctgatacaaatataaatgtacatgcacttgtacatgtcaaggggagggggggtaccaaaggtttatacattgctgattttctgttttgactcca
acttgtgcatttagttttcttgttttcgctgcagttccattcttttaaattggtatttcttttgttgtttcctttttctccatgactcttcttatt
>TRINITY_DN45638_c0_g1_i1 ZNF622
ggtaggcattcggtgtccgtcaaagttgtgcaacacgacgaaactttgagaaacgaaaggcgagtctctttgactagtaagctaaagtttgtctctttatttttctgtaaggaaagctgaaacattacggctgaaaac
aatcgatcttcgggaacaATGTCTTCCGGTACCTACACCTGTATCACATGCCGTGTGGGATTTCTGGACGGCGATCTACAGCGTGGCCATTA
TAAGACAGACTGGCATAGATATAACCTCAAACGGAAAGTAGCCGACCTGCCCCCGGTCACAGCAGATACATTCCAACAGAAAG
TGCTAGCTCAGCGGGCAAAAGTCCAGGAGCAGAGTAAAACTGAGAATGTCAAATGTGATTTATGCAGCAAACACTTCAGTACA
CAGAATGCCTACCAGAATCACCTACAGTCAAAAAAACACAAAGATGCCGTCAGTAAACAGACAGAGAAGCTAAAAACGGAAA
TCGAAAAGAAAAACGAAGAAAGTAAAACAACGGAAACTGGTTTGCCGGAAAATGAAAAGAGACTCATGAAAGACTGTGTAA
ATATGAGGCTGAAGGAAAAAATGCAAACTGCGGAATCTGATGTTGCCATGGCAACCAAAGGAAAAGGAGCAACAGGAAAAAC
AAAAATGGAAATTTCAGATGATGATGATTCTGATGCGGAGTCTGTTGACAGCTGGGATGAGAACTCTCTAGGAATAGAGGAGTG
TTTGTTCTGTTCACATATCAATTCAAGCATGGAGAAAAATATCGAACATATGACAGTCAAACATAGTTTCTTCCTCCCAGACGCA
GAATATATCACAGATTTGGAGGGTCTTATTTGTTATCTAGGAGAGAAGATAGGAGTGGGACATGTGTGTTTATGGTGTAACGAGA
AAGGGAAAACATTTCACAGTACAAAGGCTGTACAGAAACACATGCTCGACAAAGGTCACTGTAAGATACTACATGAGGGGGAT
ACCGTATATGAATTTGCTGATTTCTATGACTATAGGAGCAGTTATCCTGACTACCAGGAAGGCACAGAACAGGATTCTGGGAATG
ACGACCCTGAGAGGGATGTAGAGATGGAGGATGACAGCGATGATGAAGTCACGGAGCAGGAAATCAAGGGAGACGGTTACGA
ACTAGTGCTACCATCTGGTGCCACCATTGGACACAGATCATTGATGAAGTACTATCGGCAGAATCTACCGAACAGATCTTACGAT

AAAACAAAGACGATACTACCTAAGATGCTGGCACAGTATAAGGCACTGGGATGGACCGGCACCACAGGCGTGGTAGCTCAGAGACGTGCCAAGGACCTCGGCTACATGCAGAGATTAAAGTCACGGAAATACATGGATGTGGGCATTAAGGCAAATAAGCTACAGACACACTTCAGACCTCAAGTCATCTTCTAAgctacgactaaagattaaaagaagcattttttttggggggggggttgccagtaaaatttttgaagagggaatcaatgcaactatatttggaacagtcatggaaatgaaattatgcctggattatttattttgtgtacctaAAAACAAATTTTCTTTGAATTTAAAGTGttttgtgtttgaaagtatttgatagaggatctgttttcgtattcagacacccatgtttgattacatttattgaagttgtactaaaactttttacttagcaagacacttatgattataatcaaattgaaaaatgtacaggcatataaagaatggggcagaaacagatgcatattttaaatttctatctattttgatgattcttatgtgaacaagatggaattactggtaacagaatgaataatattgtgcaaataataattacactcatgggggaagatattgtcatcattgctggggtgattctatcatacaaaatggtgaataaattgacctaaatctaaaaaaaaa
